# Endogenous tracking reveals paralog-specific targets for phosphatidic acid generation during phospholipase C signaling in living cells

**DOI:** 10.64898/2026.03.02.709100

**Authors:** Claire C. Weckerly, Olivia L. Murtagh, Tiernan Swayhoover, Joshua G. Pemberton, Ku-Lung Hsu, Gerald R. V. Hammond

## Abstract

Phosphatidic acid (PA) is an essential intermediate generated during phospholipase C (PLC) signaling, but its regulation is complex. PA can be generated by ten different diacylglycerol kinase paralogs (DGKs) and two different phospholipase D paralogs (PLDs) in mammals. Because these enzymes are activated under diverse conditions and at various membranes, understanding paralog-specific contributions to PA production is critical for therapeutic development of drugs that modulate the PLC pathway. To address this, we aimed to characterize the paralog specificity of the DGK inhibitors R59022 and BMS-502 against individual endogenously-tagged DGK paralogs in live cells. We found that R59022 and BMS-502 both recruited endogenous DGKα to the plasma membrane, and inhibited the catalytic fragment of DGKα when ectopically localized to the mitochondrial outer membrane. However, at its effective dose, R59022 paradoxically increased PA levels, while BMS-502 functioned as a potent inhibitor. Live-cell imaging experiments using BMS-502 with carbachol stimulation of endogenous muscarinic receptors showed that inhibition of both DGKα and the PLDs is needed to substantially reduce PA levels during PLC activation. Our findings both identify paralog-specific druggable targets for modulating PLC signaling events, and establish a new platform for characterizing DGK and PLD activity in living cells.

## Introduction

Phosphatidic acid (PA) is a key signaling and metabolic lipid. After phospholipase C (PLC) cleaves phosphatidylinositol 4,5-bisphosphate (PIP_2_), into diacylglycerol (DAG) and inositol (1,4,5)-trisphosphate (IP_3_), PA is produced downstream. The PA then plays a critical role in replenishing PIP_2_ levels and maintaining plasma membrane (PM) homeostasis. PLC activation by G-protein–coupled receptors generates PA through two pathways: diacylglycerol (DAG) is phosphorylated by diacylglycerol kinases (DGKs), or protein kinase C (PKC) activation by DAG and elevated Ca^2+^ levels stimulates phospholipase D (PLD) to hydrolyze phosphatidylcholine (PC) and make PA (1). Experimental evidence shows that both DGKs and PLDs do contribute to PA levels (2–4), but the roles of specific paralogs in this pathway remain unclear. Defining these roles is essential to guide targeted therapies for PLC-related diseases such as hypertension or PA/DAG-mediated signaling such as that in T cell proliferation (5–10).

Mammals express ten DGK paralogs (α, β, γ, δ, ε, η, ι, κ, θ, ζ) grouped into five subtypes (Type I-V) based on their regulatory domains, as well as two PLD paralogs (PLD1 and PLD2) (11, 12). One common regulatory mechanism for the DGKs and PLDs is PM translocation, as has been demonstrated for DGKα, DGKδ, DGKγ, DGKθ, and DGKζ (13–19). PLD1, which is typically Golgi-associated, can also translocate to the PM, and has been implicated in PA production during muscarinic receptor signaling (3, 20–22). On the other hand, PLD2 resides at the PM even at rest (21, 23, 24). However, context matters; for example, DGKα localizes to the PM during T cell receptor activation, but moves to the perinuclear region upon IL-2 stimulation of lymphocytes (14, 25). Thus, as cellular localization of these enzymes is highly dependent on the context and specific paralog, it is important to screen each individual paralog for its membrane association under various signaling contexts.

To distinguish between DGK activity and PLD activity in the production of PA, we turned to the widely used DGK inhibitor 6-{2-{4-[(4-fluorophenyl)phenylmethylene]-1-piperidinyl}ethyl}-7-methyl-5H-thiazolo-(3,2-a)pyrimidin5-one, or R59022. Its reported IC_50_ ranges from 2.8–6.7 *μ*M in red blood cell membranes or intact platelets (26, 27) to 25.4 *μ*M when using DGKα lysates in liposome assays (28).

The paralog-specificity of this inhibitor has been debated. R59022 inhibited DGKα in early studies and was less effective against other paralogs (25, 29, 30). In agreement with this, recent work showed that R59022 only inhibited DGKα and not DGKι (28). In a full panel of DGKs, however, R59022 showed activity against DGKα, DGKε, and DGKθ (31).

R59022 has been used across many cell types (2, 25, 32–47) where it was shown to effectively increase DAG, decrease PA, and modulate signaling events dependent on these lipids. R59022 even reduced glioblastoma/melanoma tumor growth *in vivo* (48). Importantly, its effects are often stimulus-dependent and R59022 alone produced no phenotype (2, 27, 43, 44, 49).

Interestingly, in several studies, higher doses of R59022 or long incubation times with the inhibitor would paradoxically increase PA, induce morphological changes, or begin to inhibit effects that were potentiated by lower levels of R59022 (27, 35, 43, 49, 50). Despite these reports, many of the aforementioned studies used R59022 at high levels and did not describe any unusual effects. Of particular interest to us, studies using a high concentration of R59022 or long incubation periods while imaging lipid biosensors still observed the expected decreases in the PM localization of PA biosensors and increases in the PM localization of DAG biosensors (51–54). However, in our hands, we saw cytotoxic effects of R59022 and an increase in PA levels using our PILS-Nir1 biosensor for PA **(Fig. 1)**, so we aimed to characterize the paralog specificity, IC_50_, and cellular effects of R59022 in our own HEK293A cells.

**Figure 1.**
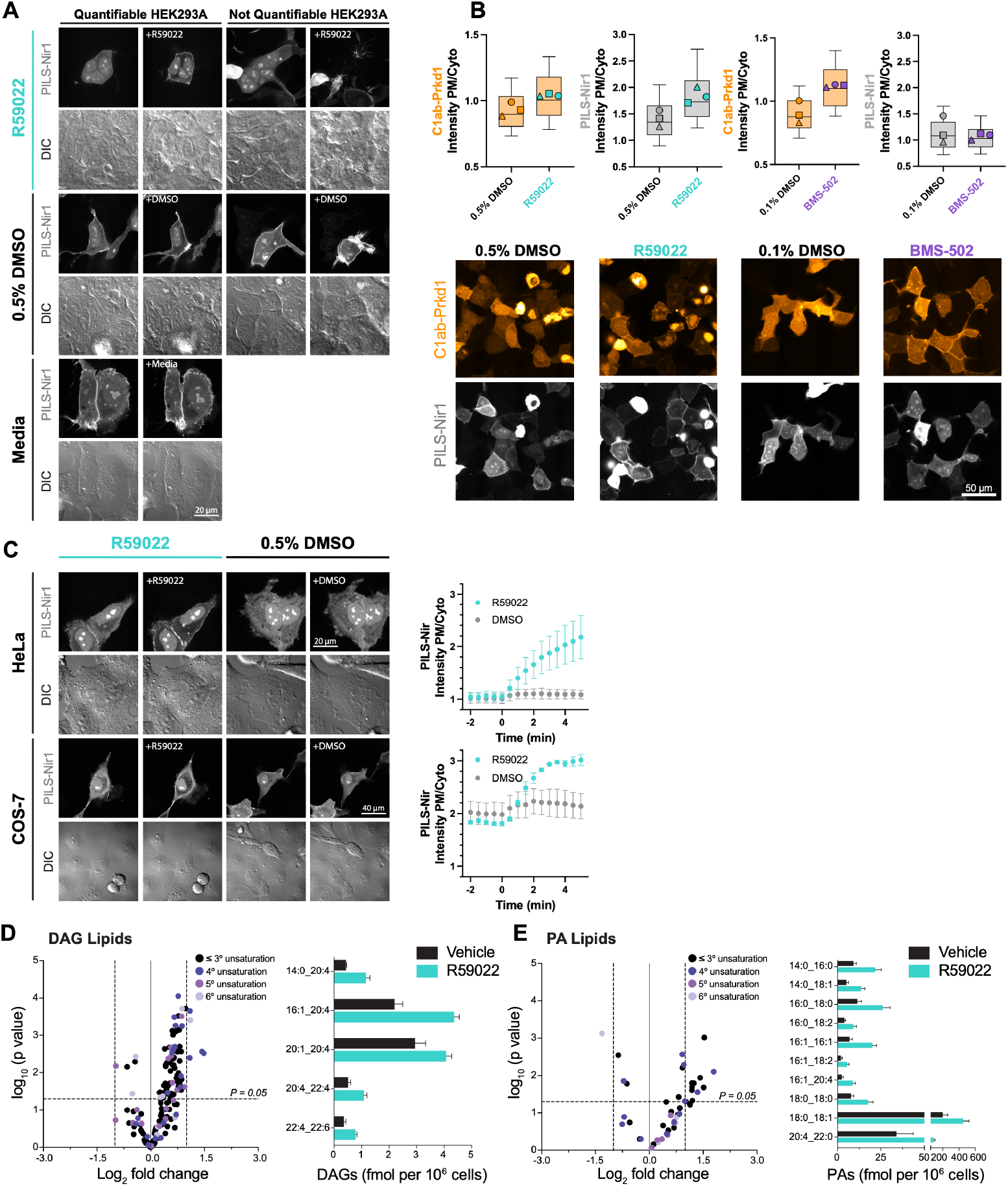
R59022 is toxic to cells and increases PA levels. **(A)** Representative images of acute stimulation of HEK293A cells with 50 *μ*M R59022, 0.5% DMSO, or cell imaging media. The PILS-Nir1 PA biosensor is shown to track PA production at the PM, and the differential interference contrast (DIC) is shown to demonstrate cell morphology. Stimulation with R59022 or 0.5% DMSO resulted in a majority of cells being not quantifiable (right). Cells stimulated with imaging media were all quantifiable (left). **(B)** Cells were treated with 50 *μ*M R59022, 1 *μ*M BMS-502, or their respective DMSO controls for 1 hour before large images of the field of cells were taken using confocal microscopy. Representative images are shown. The PM localization of the DAG biosensor C1ab-Prkd1 and PA biosensor PILS-Nir1 was quantified as the ratio of fluorescence intensity at the PM to that within the cytoplasm. The box and whisker plot shows the pooled data of 294-374 cells. The box shows the interquartile range and the whiskers show 10-90^th^ percentile. The large symbols denote the means of the individual experiments, with data from the same replicate matched by symbol shape. **(C)** Representative images of acute stimulation of HeLa and COS-7 cells with 50 *μ*M R59022 or 0.5% DMSO. Graphs show quantification of PILS-Nir1 at the plasma membrane, showing grand means ± SEM of 3 experiments (analyzing 7-15 cells per replicate experiment). **(D)** Targeted lipidomics show statistically significant upregulation of unsaturated DAGs in HEK293A cells treated with R59022 (n = 3 replicates). The horizontal dashed line in the volcano plot indicates the statistical significance cutoff of p = 0.05. The vertical dashed line shows the fold change cutoff of R59022/DMSO comparisons (Log_2_FC = 1). Statistical significance was determined using a Holm-Sidak correction following an unpaired t-test. The bar graph shows the DAG species that were significantly upregulated. Data is presented as the mean ± SEM for the 3 replicates. **(D)** As in C, but showing that saturated PA species were statistically significantly upregulated with R59022 treatment.

To help us understand the effects of R59022, we compared it to the newly characterized DGK inhibitor, BMS-502. BMS-502 was discovered in a phenotypic screen for compounds that could serve as a combined therapy with PD-1 checkpoint inhibitors in T cells. It potently and specifically inhibited DGKα, DGKζ, and DGKι at nanomolar levels (55, 56). Interestingly, it also induced DGKα and DGKζ translocation to the PM in Jurkat cells, a mechanism requiring further study as PM translocation of DGKs tends to be associated with enzyme activation, not inhibition (56). Still, this inhibitor is extremely promising since when combined with PD-1 antibodies, BMS-502 nearly eliminated tumor growth in mice (56).

The aim of this study was to characterize R59022 and BMS-502 within the native cellular environment to try and get a more robust understanding of their cellular effects, so that we could use these inhibitors to deconvolve DGK and PLD contributions to PA levels. Here, we used live-cell imaging of individual DGK paralogs to compare effects of R59022 and BMS-502, as these cellular assays were recently used to effectively characterize a novel DGK inhibitor (57). We tracked PM localization of the endogenous enzymes and directly tested inhibitor activity on DGKα within live cells, allowing us to determine paralog specificity, EC_50_ values, and IC_50_ values as one would traditionally do with purified enzymes and synthetic substrates *in vitro*. Our results show that BMS-502 is a potent DGKα, DGKθ, and DGKζ inhibitor, while R59022 has significant off-target effects, most notably the induction of a paradoxical *increase* in cellular PA. We then use BMS-502 to identify DGKα and PLD2 as key contributors to PA generation during PLC activation. Overall, this work highlights new therapeutic targets and introduces a versatile live-cell platform for probing DGK regulation and inhibition.

## Results

### R59022 is toxic to cells and increases PA levels

To start testing the efficacy of R59022 against PA production, we turned to our newly created PA biosensor, PILS-Nir1 (4). This biosensor allows us to quantify relative PA levels by determining the ratio of sensor fluorescence intensity at the PM to that within the cytosol (cyto). Thus, a stimulation that induces PA production at the PM will cause the sensor to translocate to the PM and bind the PA, which results in a higher ratio of PILS-Nir1 fluorescence PM/cyto.

After imaging HEK293A cells expressing the biosensor for a 2-minute baseline, we acutely added 50 *μ*M of R59022 to see if this high dose had the reported off-target effects in our cells. Interestingly, we saw that upon addition of the drug, many cells blebbed, shrunk, rounded up by coming off the dish or otherwise washed away **(Fig. 1A)**. These cells were labeled as “not quantifiable cells” due to the lack of a consistent area of the cytoplasm where a region of interest could be drawn during analysis. The differential interference contrast (DIC) image of these cells shows high confluence of the cultures, indicating that the cells didn’t detach due to lack of cellular contacts. It also confirms that significant changes in cellular morphology occurred. Only a few cells remained where the cytosol and PM could be monitored throughout the entire time course, and we deemed these cells to be “quantifiable cells”. To our surprise, these cells showed an increase in PA levels upon R59022 addition, seen by the increased PM labelling of PILS-Nir1 **(Fig. 1A)**.

As a control, we added 0.5% DMSO to the cells, which is an equal volume of DMSO to the volume of R59022 used previously. Surprisingly, this DMSO vehicle also led to a majority of the cells being “not quantifiable”, and those that we could analyze also showed an increase in PA levels **(Fig. 1A)**. To determine if the increase in PA was due to mechanical stimulation of the cells while pipetting the solutions on, we “stimulated” the cells with cell imaging media. These cells were all considered quantifiable and only showed a very subtle increase in PA levels **(Fig. 1A)**. R59022 used requires levels of DMSO that are cytotoxic to cells.

As acute stimulation with these levels of R59022 and DMSO seemed too harsh on the cells, we turned to long-term stimulation instead. This experimental set up allowed us to slowly pipette the solutions on to the cells. We then let the cells acclimate to the stimulation for 1 hour before using confocal microscopy to take an image of a large field of cells. This gentler application appeared to improve the overall morphology of the culture and thus reduce cytotoxicity. Using the C1ab-Prkd1 biosensor for DAG levels, we confirmed that 50 *μ*M R59022 caused the expected accumulation of DAG versus the 0.5% DMSO vehicle control **(Fig. 1B)**. Interestingly, we saw that R59022 also elevated PA levels as compared to the control, seemingly agreeing with our observations in the acute experiment **(Fig. 1B)**. This experiment suggests that R59022 has effects inherent to the compound that are not just attributable to the high concentration of DMSO that it is dissolved in.

To investigate whether the acute toxicity and induced PA increases were unique to HEK293A cells, we repeated these initial experiments with HeLa and COS-7 cells. Unlike the HEK293A cells, application of 50 *μ*M R59022 or 0.5% DMSO did not lead to obvious changes in cellular morphology or acute signs of toxicity in either cell line **(Fig. 1C)**. Therefore, toxicity seems to be a unique feature of HEK293A cells. However, R59022 induced a robust increase in PILS-Nir1 at the PM, suggesting a consistent and paradoxical increase in PA levels induced by this compound **(Fig. 1C)**.

We then wanted to see if this same elevation in PA happened with BMS-502 stimulation. When 1 *μ*M BMS-502 was used, we confirmed the anticipated increase in DAG, but saw some inhibition of PA levels compared to a 0.1% DMSO control **(Fig. 1B)**. This suggests that BMS-502 may avoid the unusual PA increase seen with R59022.

To confirm the increase in PA seen with R59022, we turned to low-microflow targeted lipidomics (58) to look at DAG and PA species. Indeed, this method showed an increase in several unsaturated DAGs and PAs **(Fig. 1D,E)**. However, the acyl chains of the DAG species elevated by R59022 are not the acyl chains that are typically thought of as DGKα-specific substrates, suggesting that R59022 is more of a general DGK inhibitor (59). Thus, it appears that R59022 broadly interacts with the DGKs in a way that results in elevated PA.

Overall, we saw that 50 *μ*M R59022 was cytotoxic when acutely added to HEK293A cells, but otherwise caused an increase in PA levels in three cell lines. This effect is counter to the expected effects of a DGK inhibitor, and can lead to the re-interpretation of many existing experiments that employed it. High levels (1 *μ*M) of BMS-502 did not show these same effects in our imaging data. Thus, we set out to characterize the dose-response of R59022 and BMS-502 in order to determine an effective dose that can be used for our live-cell imaging experiments.

### R59022 and BMS-502 recruit endogenous DGKs to the PM

We first wanted to look at the effects of R59022 and BMS-502 on specific DGK paralogs using our new *in cellulo* biochemistry approach. To do this, we turned to our recently described HEK293A cell lines where CRISPR/Cas9 was used to introduce a fluorophore onto the endogenous gene loci of the DGK paralogs (57). These cell lines each express one of the DGK paralogs: DGKα, DGKδ, DGKε, DGKη, DGKθ, or DGKζ with a fluorescent tag (either NeonGreen (NG) or StayGold (SG)) at endogenous levels. The selected paralogs are the only 6 DGK paralogs found in HEK293A cells (of 10 total mammalian paralogs) (60). Notably however, these 6 paralogs span each of the 5 subtypes of DGKs, so we anticipate these tagged cell lines will provide broad insights into how the DGK paralogs are regulated.

Using these cell lines, we first wanted to know how the different paralogs respond to R59022 or BMS-502, especially as membrane translocation of DGKs was previously used to show the paralog specificity of BMS-502 (56). To do this, we imaged the cells using total internal reflection fluorescence microscopy (TIRF), a technique that selectively excites fluorophores on or near the PM. TIRF removes much of the cytosolic noise, allowing for single molecules to be imaged. We stimulated each cell line with either 50 *μ*M R59022 or 1 *μ*M BMS-502 for 5 minutes while imaging. As noted in **Fig. 1A**, 50 *μ*M R59022 is toxic and induces extensive rounding in many cells. However, imaging the cell footprint in TIRF and analyzing a relatively small 10×10 *μ*m region allowed much more consistent tracking of cells than seen in confocal. Interestingly, we saw that R59022 caused large PM recruitment of NG-DGKα and SG-DGKζ, and modestly recruited DGKθ-SG. This is seen by an increase in the number of fluorescent particles within the cell membrane footprint in TIRF images (**Fig. 2C**) and was quantified as the number of particles found in a representative 100 *μ*m^2^ area (**Fig. 2A**). Addition of BMS-502 showed a similar pattern of NG-DGKα, DGKθ-SG, and SG-DGKζ recruitment to the PM (**Fig. 2B,C**). Notably, the magnitude of NG-DGKα recruitment was less with BMS-502 than it was with R59022.

**Figure 2.**
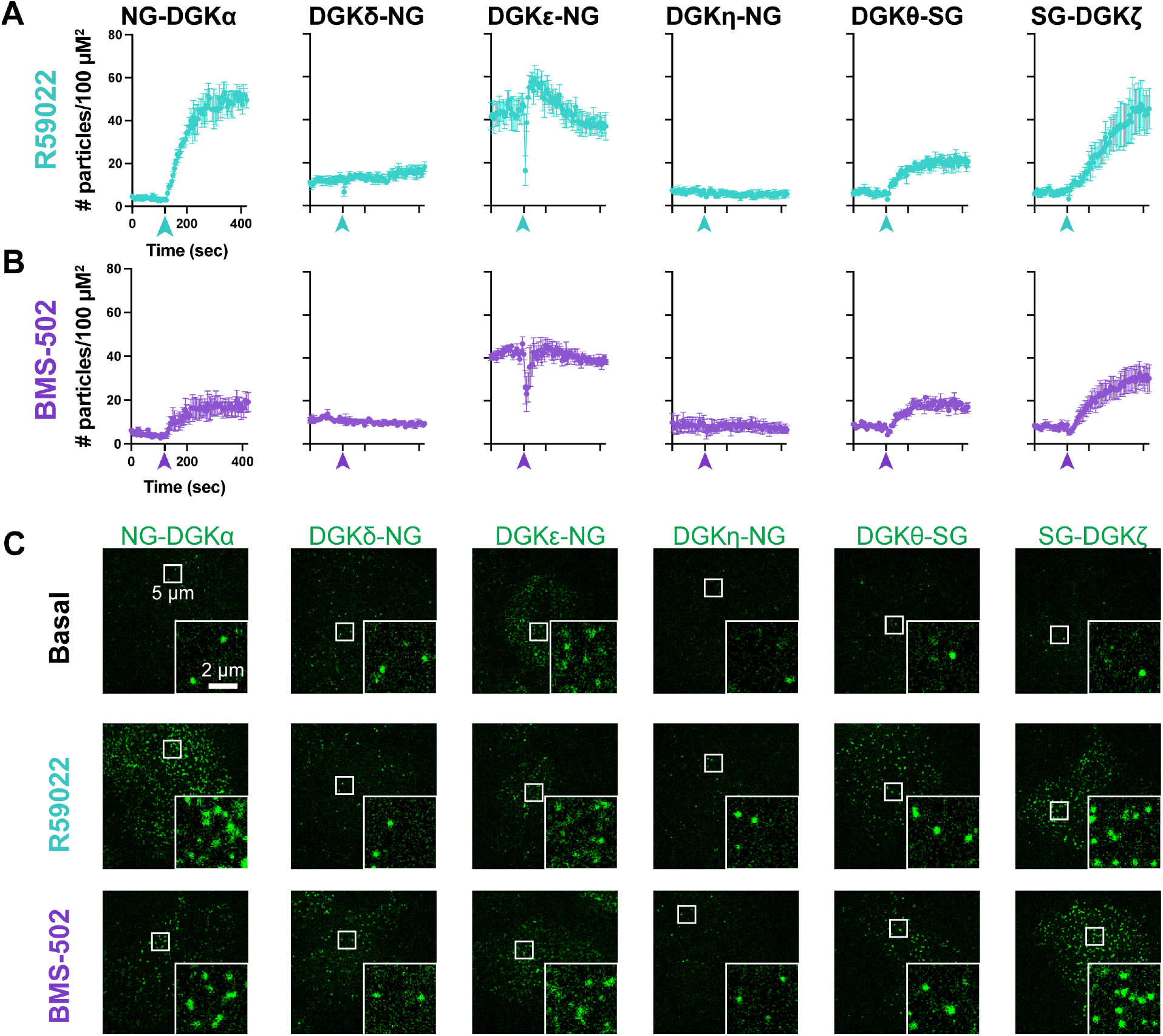
R59022 and BMS-502 recruit endogenous DGKs to the PM. **(A)** 50 *μ*M R59022 was added to the different tagged cell lines as indicated by the arrow. Recruitment of the DGK particles to the PM was determined using TIRF microscopy to quantify the number of particles present in a representative 100 *μ*m^2^ area of the cellular footprint. **(B)** As in A, but using 1 *μ*M BMS-502. Data shown is the grand means of 3-5 experiments ± SEM. A total of 9-18 cells were analyzed. **(C)** Representative TIRF images of A and B. The basal cell shown is the same cell shown after R59022 stimulation, which is representative across the different stimulations used.

The BMS-502-induced DGK translocation agreed with previously published data (56), however PM translocation caused by R59022 was unprecedented. In fact, R59022 has been shown to inhibit DGKα translocation to the PM in lymphocytes (25). Although, the structurally similar DGK inhibitor R59949 did prolong DGKα association with the PM after CCh-induced translocation (15, 26). Therefore, we wanted to study the R59022-induced translocation of DGKα further.

We saw an increase in PA levels with R59022 but not BMS-502, and we saw that R59022 caused more NG-DGKα translocation than BMS-502 did. Therefore, we hypothesized that R59022 recruitment of NG-DGKα to its PM-localized substrate may overcome the inhibitory effects of R59022 on catalytic activity. In essence, we hypothesized that R59022 was acting as a competitive inhibitor. In a competitive inhibition model, increasing the DAG concentration around NG-DGKα by bringing it to the PM would cause DAG to outcompete R59022 for binding of NG-DGKα, thereby increasing PA levels. To this end, we devised an *in cellulo* biochemistry method to compare R59022’s ability to recruit NG-DGKα with its ability to inhibit PA production.

### R59022 recruits DGKα to the PM and inhibits its catalytic activity in the micromolar range

If R59022 increases PM localization so that the substrate availability outcompetes inhibition by R59022, then we would expect that the effective concentration of R59022 needed to recruit NG-DGKα would be significantly lower than the inhibitory concentration needed to inhibit the phosphorylation reaction.

To determine the effective concentration of R59022 for NG-DGKα recruitment, we repeated the TIRF experiment as described above, but with gradually higher doses of R59022. Here we saw that increasing doses of R59022 correlated with increased NG-DGKα recruitment to the PM **(Fig. 3A)**. Interestingly, we also noted that the 1.5% DMSO control used here caused a small and transient recruitment of NG-DGKα to the PM, which could explain some of the increases in PA levels we had seen previously. Note that higher doses of R59022 were tolerated in these TIRF experiments, as only a 100 *μ*m^2^ area of the cell membrane is analyzed, so even if the cell blebs or shrinks upon stimulation, an area large enough for analysis usually remains.

**Figure 3.**
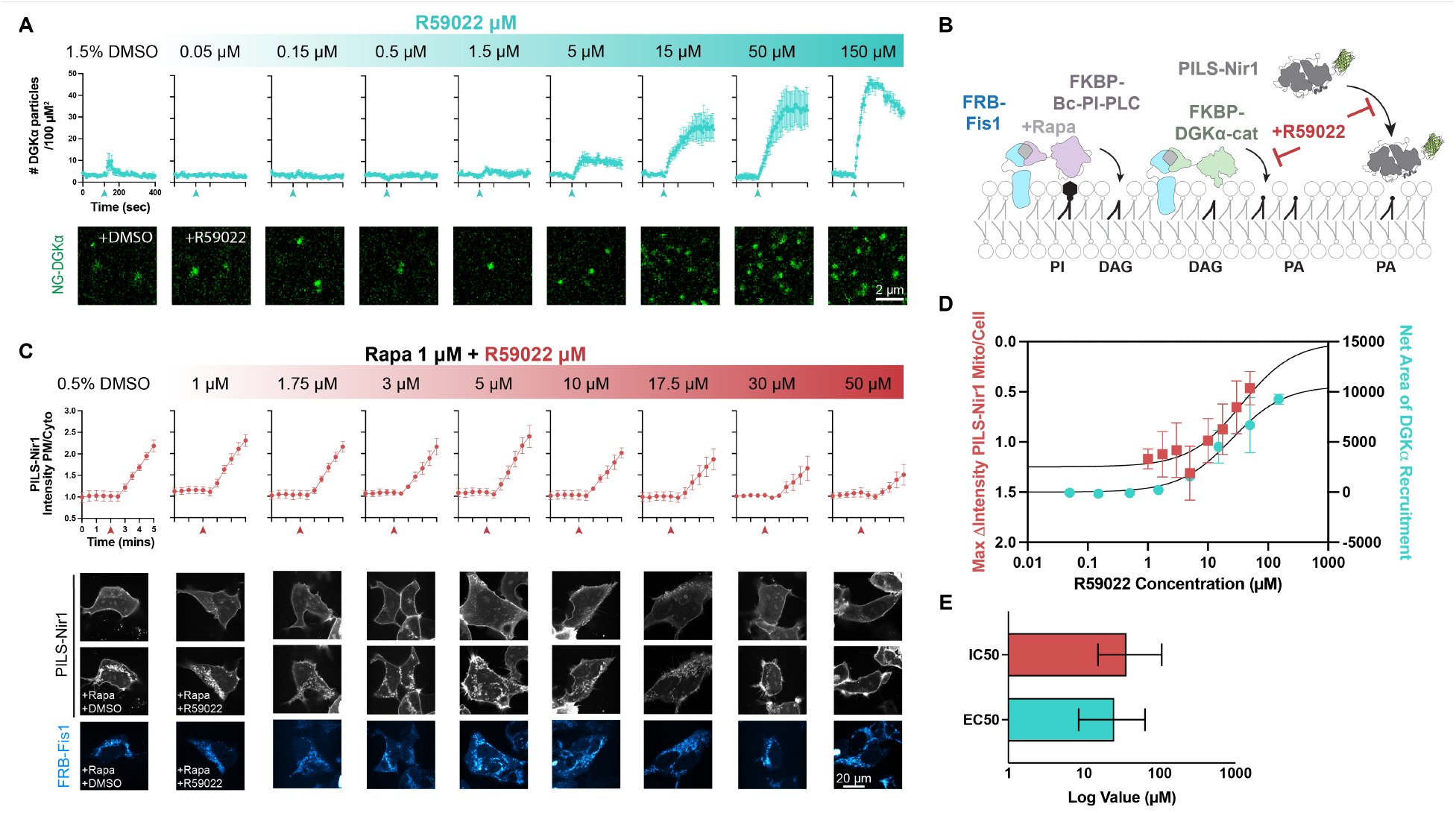
R59022 recruits DGKα to the PM and inhibits its catalytic activity in the micromolar range. **(A)** Increasing doses of R59022 were added to the NG-DGKα cells, as shown by the arrow, to determine recruitment of DGKα to the PM. Recruitment was quantified as the number of particles present in a representative 100 *μ*m^2^ area of the cell. Representative TIRF images after drug addition are shown below. Data shown is the grand mean of 3 experiments ± SEM. A total of 10-13 cells were analyzed. **(B)** Schematic of the chemically inducible dimerization system used to produce PA at mitochondria. **(C)** Quantification of the intensity of PILS-Nir1 at the mitochondria to that within the whole cell when various doses of R59022 were used to inhibit the chemically inducible dimerization system (which was activated by a constant dose of rapamycin). Cells were stimulated as shown by the arrow. The grand means of 3 experiments ± SEM is shown; a total of 32-44 cells were analyzed. Representative confocal images of PILS-Nir1 before and after stimulation, and of the mitochondrial marker FRB-Fis1 after stimulation are shown. **(D)** The data from A and C was re-plotted as a dose response curve. To look at inhibitory effects, the max change in PILS-Nir1 intensity at the mitochondria was plotted using an [inhibitor] vs response (three parameters) least squares fit. The data is shown on an inverse y-axis to more easily compare the inhibition curve to the DGKα recruitment curve. The bottom was constrained to 0, and the IC_50_ was constrained to > 0. Degrees of freedom = 22. R^2^ = 0.3589. To look at DGKα recruitment, the net area under the curve of the data in A was plotted using [agonist] vs response (three parameters) least squares fit. The bottom was constrained to 0, and the EC_50_ was constrained to > 0. Degrees of freedom = 22. R^2^ = 0.9211. **(E)** A bar graph showing the IC_50_ and EC_50_ values calculated above with the whiskers showing the 95% confidence interval.

To determine the concentration of R59022 needed to reduce DGKα activity, we turned to a chemically-inducible dimerization system that produces PA on mitochondria **(Fig. 3B)** (4). We co-expressed a phosphatidylinositol-specific PLC from *Bacillus cereus* (FKBP-*Bc*PI-PLC) and the catalytic domain of DGKα (DGKA paralog b residues 394-773; referred to as DGKα-cat), both with an added FK506-binding protein (FKBP) from the mTOR complex. Then, the FKBP-rapamycin-binding domain (FRB) from mTOR was targeted to the mitochondria by the addition of the localization sequence from the mitochondrial fission 1 protein (FRB-Fis1). When we stimulated the cells with rapamycin, the FKBP and FRB-Fis1 dimerized, localizing the FKBP-*Bc*PI-PLC and FKBP-DGKα-cat to the mitochondria. There, the FKBP-*Bc*PI-PLC cleaved PI to produce DAG. The FKBP-DGKα-cat then phosphorylated the DAG to produce PA, which we detected with the PA biosensor PILS-Nir1. We added R59022 to the system in increasing doses to inhibit FKBP-DGKα-cat activity and reduce PA production **(Fig. 3B)**. Thus, this assay resulted in a simplified system where the DGKα catalytic fragment was tethered to the mitochondrial membrane rather than its localization being affected by the presence of R59022. This system was used recently to characterize a novel DGK inhibitor (57).

We first confirmed that simultaneously adding different doses of R59022 with a constant concentration of rapamycin did not affect the ability of rapamycin to recruit the FKBP constructs to mitochondria **(Fig. S1A, B)**. Looking at the PILS-Nir1 response to PA production, we saw that increasing the dose of R59022 did reduce the amount of PA made at the mitochondria **(Fig. 3C)**.

We then plotted the data from **Fig. 3A** and **Fig. 3C** as dose-response curves to calculate the effective concentration of R59022 to cause 50% NG-DGKα recruitment (EC_50_) and the inhibitory concentration to cause 50% inhibition of PA production at the mitochondria (IC_50_). We plotted the PA inhibition curve on an inverse y-axis to more easily compare the shapes of the two curves **(Fig. 3D)**. If R59022 was more potent at recruiting DGKα than it was at inhibiting it, we would see the curve of NG-DGKα recruitment be left-shifted with a low EC_50_ value, and the inhibition curve be right-shifted with a higher IC_50_ value. Instead, we see that the dose-response curves show the same shape, indicating no difference in the potency of R59022 for these two activities **(Fig. 3D)**. This is further confirmed by comparing the IC_50_ and EC_50_ values, which are comparable and fall within a similar 95% confidence interval **(Fig. 3E)**.

Thus, our idea that R59022 is a competitive inhibitor that recruits NG-DGKα more potently than it inhibits DGKα is not supported by our data. However, the data does suggest that R59022 inhibits DGKα in the 10-50 *μ*M range, which is unfortunately when the cytotoxic effects of R59022 and/or the DMSO can also be seen within the cells (**Fig. 3C**, note the cells beginning to round up when treated with 30 *μ*M or 50 *μ*M R59022). Overall, our data does not show R59022 to be an effective DGK inhibitor for use in live HEK293A cells, so we turned to BMS-502 instead.

### BMS-502 recruits DGKα to the PM and inhibits its catalytic activity in the nanomolar range

Since we saw that BMS-502 did not raise PA levels **(Fig. 1B)** and its ability to recruit DGKα has been previously characterized (56), we wanted to find a useful concentration of BMS-502 for use in our HEK293A cells. To do this, we repeated the same experiments from **Fig. 3** now using BMS-502. We saw that BMS-502 also recruits NG-DGKα to the PM in a dose-dependent manner **(Fig. 4A)**. In our mitochondrial assay, we observed that BMS-502 did not affect the ability of rapamycin to recruit the FKBP constructs to the mitochondria **(Fig. S1C,D)**. Additionally, it inhibits FKBP-DGKα-cat at the mitochondria, with a concentration of 1000 nM (1 *μ*M) resulting in almost complete inhibition **(Fig. 4B)**. Importantly, the cells treated with this highest dose of BMS-502 still maintained normal morphology, indicating that BMS-502 at high doses does not have the same cytotoxicity that we see with R59022 **(Fig. 4B)**. We then plotted the dose-responses of NG-DGKα recruitment against FKBP-DGKα-cat inhibition and saw that the EC_50_ and IC_50_ values overlapped significantly, as we saw with R59022 **(Fig. 4C,D)**. Unlike R59022 however, BMS-502 is effective around 100 nM, a dose where cells also retained normal morphology. This suggests that BMS-502 is a more potent and less toxic DGK inhibitor.

**Figure 4.**
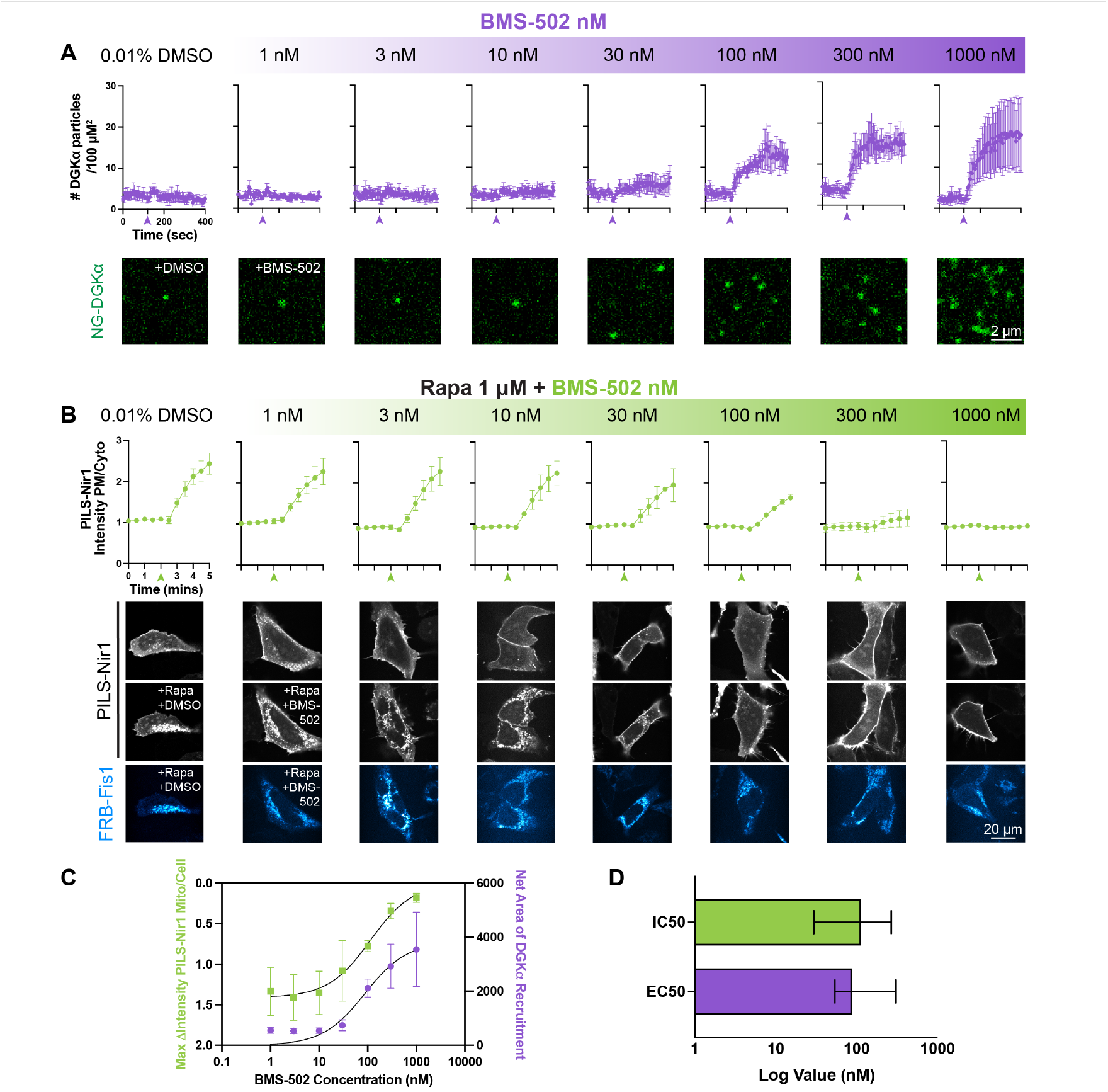
BMS-502 recruits DGKα to the PM and inhibits its catalytic activity in the nanomolar range. **(A)** Increasing doses of BMS-502 were added to the NG-DGKα cells, as shown by the arrow, to determine recruitment of DGKα to the PM. Recruitment was quantified as the number of particles present in a representative 100 *μ*m^2^ area of the cell. Representative TIRF images are shown below. Data shown is the grand mean of 3 experiments ± SEM. A total of 11-15 cells were analyzed. **(B)** As in Figure 3, a chemically inducible dimerization system was activated by 1 *μ*M rapamycin (indicated by the arrow) to produce PA at the mitochondria. Increasing doses of BMS-502 were added (indicated by the arrow) to inhibit FKBP-DGKα-cat activity, thereby causing less PILS-Nir1 (in gray) association with mitochondria, which is labelled by the FRB-Fis1 in blue. Data shown is the intensity of PILS-Nir1 at the mitochondria to that within the whole cell. The grand means of 3 experiments ± SEM is shown; a total of 30-37 cells were analyzed. Representative confocal images of PILS-Nir1 before and after stimulation, and of the mitochondrial marker FRB-Fis1 after stimulation are shown. **(C)** The data from A and B was re-plotted as a dose response curve. Inhibitory effects were quantified using an [inhibitor] vs response (three parameters) least squares fit. The data is plotted with an inverse y-axis to easier compare the inhibition curve to the DGKα recruitment curve. The bottom was constrained to 0, and the IC_50_ was constrained to > Degrees of freedom = 19. R^2^ = 0.6569. The recruitment of NG-DGKα was quantified using [agonist] vs response (three parameters) least squares fit. The bottom was constrained to 0, and the EC_50_ was constrained to > 0. Degrees of freedom = 19. R^2^ = 0.61. **(D)** A bar graph showing the IC_50_ and EC_50_ values calculated above with the whiskers showing the 95% confidence interval.

### DGKα and PLD activity are both necessary for PA production during PLC activation

Having characterized BMS-502 as an effective and safe DGKα inhibitor in HEK293A cells, we then utilized it to see how DGK activity contributes to PA levels after PLC activation. To ensure steady-state inhibition of DGK activity before receptor stimulation, cells were pre-incubated with BMS-502 for 30 min. After the pre-treatment with inhibitor, the cells were imaged for 2 minutes baseline, and then 50 *μ*M carbachol (CCh) was added to stimulate PLC through activation of the M3 muscarinic receptor. We again used the DAG biosensor C1ab-Prkd1 to confirm DGK inhibition. Indeed, cells treated with BMS-502 showed higher baseline DAG levels, and a sustained elevation of DAG after CCh addition as compared to the DMSO control **(Fig. 5A,G)**. Looking at the PA response with PILS-Nir1 however, we saw that cells pre-treated with BMS-502 only had slightly inhibited PA levels **(Fig. 5B,H)**.

**Figure 5.**
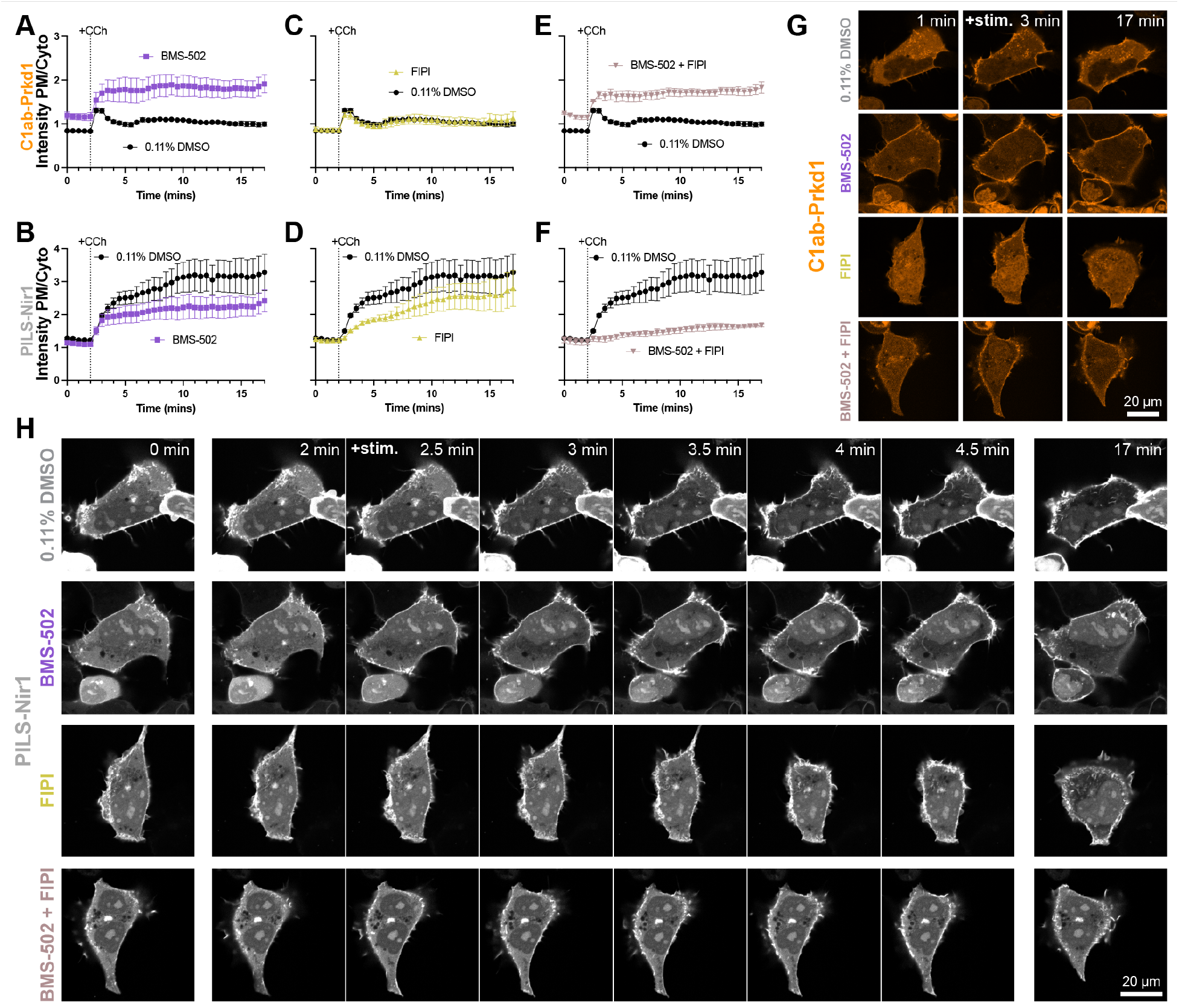
DGKα and PLD activity are both necessary for PA production during PLC activation. **(A**,**B)** 1 *μ*M BMS-502 or 0.11% DMSO as a vehicle control were added to cells for 30 minutes. The cells were then imaged on the confocal microscope for 2 minutes of baseline imaging, then 50 *μ*M CCh was added to activate PLC, as indicated by the dotted line. Association of C1ab-Prkd1, a DAG biosensor **(A)**, and PILS-Nir1, a PA biosensor **(B)**, with the PM was quantified as the ratio of intensity of sensor at the PM to that within the cytoplasm. **(C**,**D)** As in panels A and B but using 750 nM FIPI or 0.11% DMSO control (Control data is duplicated to aid in comparison between responses). **(E**,**F)** As in panel A and B but using both BMS-502 and FIPI treatment simultaneously or a 0.11% DMSO control. (Control data is duplicated between graphs to aid in comparison between responses). Data shown is the grand means of 3 experiments ± SEM; a total of 29-42 cells were analyzed. **(G)** Representative confocal images of the C1ab-Prkd1 DAG biosensor are shown, depicting the peak DAG production at 3 minutes, and the final timepoint at 17 minutes. **(H)** Representative confocal images of the PILS-Nir1 response at the first timepoint, in the first few minutes after 50 *μ*M CCh addition, and at the last timepoint after cells were pre-incubated with the respective treatments. These are the same cells as shown in G.

We suspected that BMS-502 was only somewhat effective during PLC activation because CCh will also activate the PLDs which can contribute to PA production in a DGK-independent pathway. To look at PLD activity, we used the PLD inhibitor 5-Fluoro-2-indolyl des-chlorohalopemide (FIPI), which targets both PLD1 and PLD2 (61, 62). Pre-incubating the cells with FIPI did not affect baseline DAG levels or the DAG response to CCh **(Fig. 5C,G)**, but it did reduce PA production **(Fig. 5D, H)**.

Interestingly, FIPI seemed to inhibit the initial rate of PA production, showing a slower rate of increase in the first 2-3 minutes after CCh addition **(Fig. 5D, H)**. However, by the end of the 15-minute CCh treatment, the overall levels of PA in FIPI-treated cells were comparable to the control. This is in comparison to BMS-502 treatment, which allowed PA to increase at a similar initial rate to DMSO treatment, but by the final timepoint, the PA levels were slightly lower in BMS-502-treated cells **(Fig. 5H)**. This suggests a model where PLD produces PA immediately after PLC activation, and then DGKs supply PA in later timepoints. A model where DGK inhibition does not affect initial rates of PA synthesis is consistent with previous data looking at PA production after EGF stimulation (54).

To test the idea that the PLDs and DGKs work in combination to produce PA during PLC activation, we pre-incubated cells with both BMS-502 and FIPI before stimulating them with CCh. Here, we again saw an elevation in DAG levels, due to BMS-502 inhibition of DGKs **(Fig. 5E,G)**. Notably, the combined inhibitor treatment almost completely abolished PA production and PILS-Nir1 recruitment to the PM throughout the over the entire time course **(Fig. 5F,H)**. This data supports a need for both the DGKs and the PLDs in PA production during PLC signaling.

### PLC activation recruits DGKα to the PM, while PLD2 is basally associated with the PM

We next wanted to investigate which DGK and PLD paralogs are activated by PLC signaling, as BMS-502 seemingly inhibits DGKα, DGKθ, and DGKζ, and FIPI targets both PLD paralogs. To do this, we turned to CRISPR/Cas9 to introduce fluorescent tags onto PLD1 and PLD2. We knocked fluorophores into these paralogs using a custom homology directed repair template (HDRT) that included a fluorophore sequence sandwiched between the homology arms for that specific gene.

PLD1 was tagged on the N-terminus with NeonGreen (NG). Successful editing was confirmed by an increase in size of the PCR product that resulted from two primers that flank the insertion site **(Fig. 6A)**. For PLD2, the sequence for the eleventh strand of the NeonGreen beta-barrel (NG2^11^) was incorporated into the PLD2 gene of HEK293A cells that stably express the other ten strands of NeonGreen (NG2^1-10^). Therefore, when NG2^11^-PLD2 is translated, the NG2^11^ strand reconstitutes the full beta-barrel to make a fluorescent NG molecule (NG-PLD2) (63). In the case of PLD2, the genotyping PCR used a forward primer that bound to NG2^11^ and a reverse primer that bound the PLD2 genomic DNA. Thus, this reaction resulted in the absence of a PCR product in the parental cell line, but the presence of a product in the edited cells **(Fig. 6B)**.

**Figure 6.**
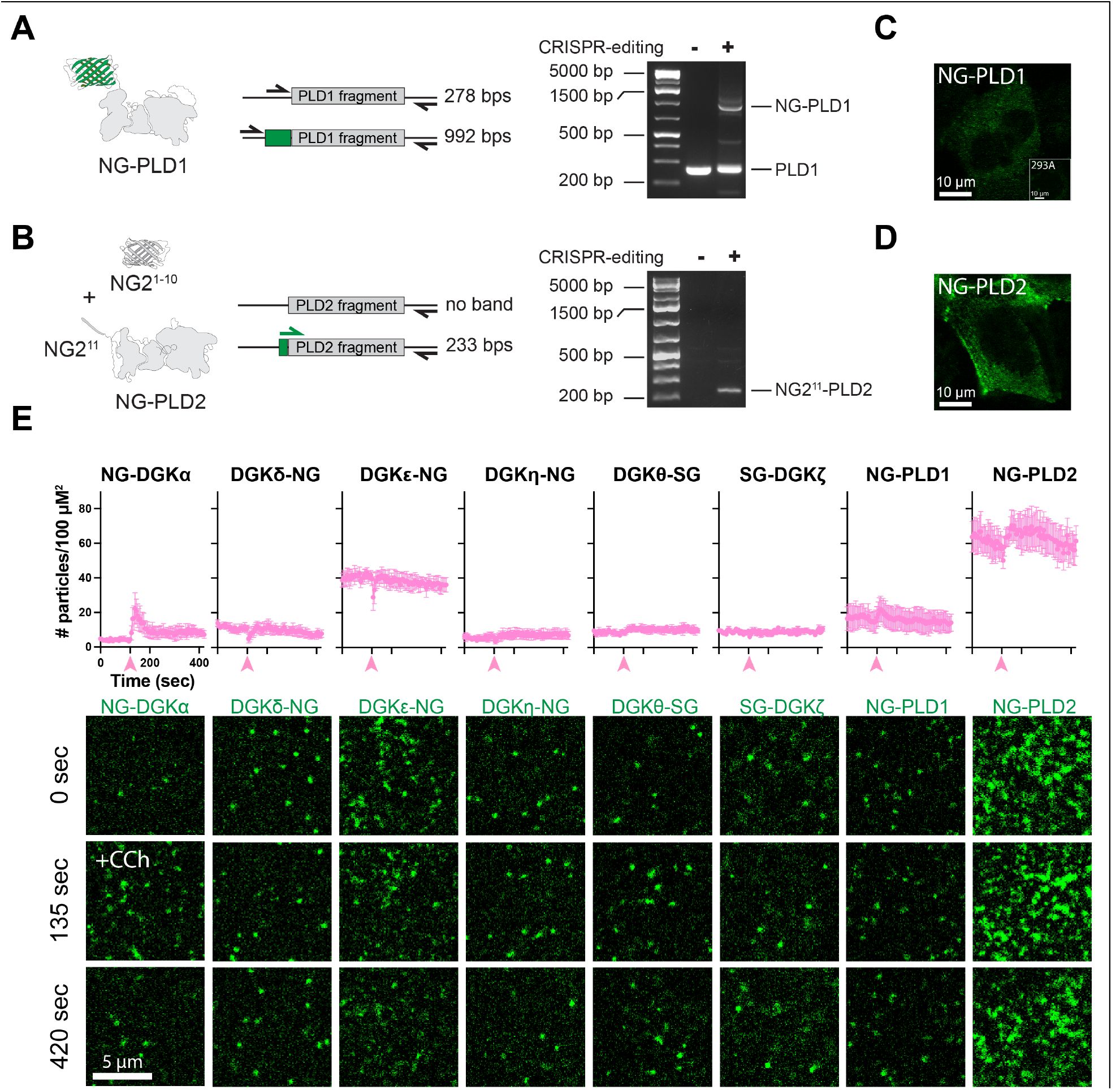
PLC activation recruits DGKα to the PM, while PLD2 is basally associated with the PM. **(A)** Cartoons showing the addition of full-length NG onto the N-terminus of PLD1 by CRISPR/Cas9 editing. Insertion was confirmed using PCR and gel electrophoresis to visualize the expected bands as shown by the diagram. **(B)** As in A, but confirming the addition of NG2^11^ onto the N-terminus of PLD2, which when translated with reconstitute the full NG barrel with NG2^1-10^. **(C)** Representative confocal image of NG-PLD1 cells show intracellular localization. Inset of parental 293A cells are shown to demonstrate that the fluorescence seen is a result of CRISPR editing and not autofluorescence. **(D)** As in C, but showing that NG-PLD2 has a plasma membrane localization. **(E)** The edited cell lines were treated with 100 *μ*M CCh to activate PLC, as indicated by the arrow. DGK and PLD recruitment to the PM was quantified as the number of particles present in a representative 100 *μ*m^2^ area of the cell. Representative TIRF images are shown below, showing the initial timepoint, the peak DGKα response at 135 sec, and the final timepoint at 420 sec. Data shown is the grand mean of 3-7 experiments ± SEM. A total of 10-18 cells were analyzed.

As PLD2 is generally considered to be localized at the PM, and PLD1 is considered to be intracellular (23), we investigated the localization of the two tagged PLD paralogs by confocal microscopy. In these images, NG-PLD1 did appear cytosolic **(Fig. 6C)** and NG-PLD2 also showed the expected PM association **(Fig. 6D)**. This confirmed that the added tags did not disrupt PLD localization within cells.

We then used another method to look at the PM localization of PLD2: we counted the particles of the DGKs and PLDs associated with the PM before any stimulation using TIRF. This can be seen in **Fig. 6E** (and the DGKs can also be seen in **Fig. 2**). We saw that before stimulation (noted by the arrow) NG-PLD2 is highly associated with the PM with about 70 particles/100 *μ*m^2^, while all the other enzymes show less than 20 particles/100 *μ*m^2^, except for DGKε-NG. While DGKε-NG shows around 40 particles/100 *μ*m^2^, it is a known ER-localized enzyme (57, 64, 65), and TIRF imaging can detect the ER structures that are in close proximity to the PM.

PLD2 is highly associated with the PM basally, but we wanted to see how the different DGK and PLD paralogs react to PLC activation by CCh. After stimulation with 100 *μ*M CCh, NG-DGKα was transiently recruited to the PM. NG-DGKα levels peaked around 135 seconds and then gradually returned to baseline. None of the other DGKs or PLDs seemed to be significantly recruited by CCh stimulation **(Fig. 6E)**. Overall, this suggests that PLC activation specifically recruits NG-DGKα to the PM, while NG-PLD2 is constitutively associated with the PM. Therefore, both of these enzymes must be inhibited, DGKα by BMS-502 and PLD2 by FIPI, to see a reduction in PA levels during PLC activation.

## Discussion

In this study, we used live cell imaging techniques backed up with lipidomic techniques to demonstrate that R59022 unexpectedly increased PA levels in cells at micromolar concentrations near its IC_50_ **(Fig. 1, 3)**. This finding potentially causes alternative interpretations for many experiments that have employed this compound. Both R59022 and BMS-502 recruited DGK paralogs, particularly DGKα to the PM, an effect which was previously reported only for BMS-502 **(Fig. 2)**. We confirmed that BMS-502 is a potent DGKα inhibitor in HEK293A cells, with an IC_50_ around 100 nM and no apparent cytotoxicity at this dose **(Fig. 4)**. Using BMS-502 with endogenously-tagged DGK and PLD paralogs, we showed that inhibition of both the DGKs and the PLDs is needed during PLC signaling to lower PA levels **(Fig. 5)**. We then showed that DGKα is specifically recruited to the PM by activation of PLC, while PLD2 shows constitutive PM localization **(Fig. 6)**.

Thus, we propose a model in which PLD2 maintains basal PA levels and contributes to early PA production after PLC activation, while DGKα produces PA during the later signaling stages, following recruitment. Overall, these findings highlight the paralog-specific contributions of the DGKs and PLDs to PA generation. Furthermore, this study utilizes fluorescently-tagged endogenous DGKs and PLDs and a mitochondrial PA assay as a platform to characterize paralog-specific drug effects and produce dose-response curves in live cells as one would with *in vitro* biochemistry.

Our work showed that the *in cellulo* assays were able to effectively characterize BMS-502 consistent with prior reports. BMS-502 is thought to directly recruit DGKα and DGKζ as part of its inhibitory mechanism, and the evidence given for this was that the IC_50_ of inhibition closely matched the EC_50_ of recruitment for each paralog. Using *in vitro* kinase assays, the IC_50_ values were 4.6-5 nM for DGKα and 2-2.1 nM for DGKζ. For PM translocation in Jurkat cells, EC_50_ values were 5 nM for DGKα and 2 nM for DGKζ (55, 56). A related small molecule inhibitor, BMS-496 was used to suggest a mechanism for DGKα and DGKζ translocation by these compounds. BMS-496 increased DGKα affinity for an arachidonoyl lipid probe without triggering Ca^2+^ release or inducing DGKα phosphorylation, indicating that the BMS compounds directly increase the affinity of DGKα for lipid substrates which could induce membrane localization (56).

Our results align with this model as our reported BMS-502 IC_50_ and EC_50_ values for DGKα were also very similar to each other **(Fig. 4)**, though higher than those reported, likely due to the different cell systems used, overexpression of FKBP-DGKα-cat, or the higher cellular substrate concentrations than those used *in vitro*. Similar to the higher concentrations needed in HEK293A cells, assays of BMS-502 in human whole-blood found that the EC_50_ of BMS-502 needed to cause elevated IFNγ was 280 nM (55). This highlights the importance of characterizing drugs within a cellular environment.

We also confirmed the reported specificity of BMS-502 for DGKα and DGKζ, but additionally found that BMS-502 recruited DGKθ (possibly indirectly, via increased DAG levels engaging its C1 domains). BMS-502 also was shown to inhibit DGKι, but since this paralog is not expressed in HEK293A cells, we did not consider it further (56). Compounds related to BMS-502 reinforce the idea that the paralogs which are inhibited by the BMS drug family are also those specifically recruited to membranes. For example, in the drug family, BMS-684 only inhibits *and* recruits DGKα, whereas BMS-118, does not inhibit *or* translocate DGKα or DGKζ (56). Altogether, this suggests that BMS-502 also likely inhibits DGKθ in HEK293A cells, as we saw DGKθ recruitment with BMS-502. However, there is a need for future studies on the interaction of BMS-502 with DGKθ.

As for R59022, the most confounding data we obtained is that both R59022 and BMS-502 recruit DGKα to the PM **(Figs. 2, 3, 4)**, yet only R59022 elevated PA levels **(Fig. 1)**. Furthermore, our results showed that R59022, like BMS-502, recruited DGKα to the PM and inhibited FKBP-DGKα-cat at the mitochondria with similar potency **(Fig. 3)**. Thus, R59022 is not acting as a simple competitive inhibitor, and increasing the DAG concentration by translocating the DGKs to the PM does not seem to be enough to overcome R59022 inhibition. This is consistent with studies of R59022 and the structurally related compound ritanserin.

Recent chemoproteomic studies using covalent probes that bind to “ligandable” residues on proteins along with analysis of the AlphaFold predicted structure for DGKα have provided insights into the structure of the DGKα active site. The catalytic domain of DGKα lies in the C-terminal end and is split into a catalytic (DAGKc) and accessory (DAGKa) domain. In the N-terminal region, DGKα contains two cysteine-rich zinc finger domains, C1a and C1b, which are similar to DAG-binding domains in PKC and help DGKα recognize substrate. The chemoproteomic studies suggest the DGKα active site spans both the DAGKa and DAGKc domains, with an allosteric site in the C1a domain that is required for activity but does not bind ATP (59, 66–68). Ritanserin binds both the C1a domain and DAGKa domain and increases the affinity of DGKα for ATP (28, 66). This data supports a mixed inhibition model where ritanserin (and likely R59022) could bind to DGKα independently through the C1a domain but have higher affinity for the DGKα-ATP complex through binding at the DAGKa domain.

Interestingly, although ritanserin was reported as being DGKα-specific, we saw R59022 recruit DGKα, DGKθ, and DGKζ **(Fig. 2)**. Of all the DGK paralogs, DGKθ and DGKζ had ligandable residues in the C1a domain similar to DGKα (66). This suggests that the C1a domain, or an active site that utilizes the C1a domain may be the source of the paralog specificity for R59022. However, additional binding sites, such as ligandable residues in the RVH domain of DGKα, may also contribute to ritanserin or R59022 binding (67). But this cannot be confirmed without further structural studies.

The presence of a ritanserin-binding site in the C1a domain highlights limitations of our mitochondrial assay, which uses the isolated catalytic domain of DGKα (residues 394-773) and thus overlooks any contributions of regulatory domains to DGKα inhibition. Moreover, the mitochondrial membrane may not fully reflect the endogenous behavior of DGKα at the PM. Interestingly, our R59022 IC_50_ value of 36.3 *μ*M (**Fig. 3**) approximates the reported *in vitro* IC_50_ of 25.4 *μ*M (28). This suggests that R59022 can inhibit DGKα when the enzyme lacks regulatory domains (as in our FKBP-DGKα-cat assay) or when DGKα regulators are not present (as in *in vitro* assays). But this inhibition does not seem to hold true when it comes to the cellular effects of R59022, likely due to the complex regulation of DGKα within the cell. Overall, we see that R59022 *can* inhibit DGKα, but *doesn’t* inhibit it in its native environment.

How then does R59022 treatment result in elevated PA levels? It is tempting to suggest that R59022 binding at the C1a domain of DGKα directly induces PM recruitment and activation. As a lipophilic drug, R59022 can incorporate into liposomes (28, 29), and thus could potentially incorporate into the PM. However, as DGKα contains an atypical C1a domain, the DAG analog phorbol 12-myristate 13-acetate (PMA) is not sufficient for DGKα PM recruitment, suggesting that R59022 binding the C1a domain would not be sufficient either (13, 69, 70).

The presence of atypical C1a domains in DGKα also imply that the increase in DAG during R59022 treatment is not sufficient to recruit DGKα to the PM. Further supporting this idea, neither ritanserin nor R59022 alter DGKα’s DAG affinity *in vitro* (28). Additionally, R59022 treatment elevated DAG species with acyl chains that are not typically selected by the DGKα C1a domain (**Fig. 1**, (59), suggesting that the increase in DAG by R59022 does not directly recruit or activate DGKα.

Alternatively, R59022 may activate PKC. R59022 has been shown to prolong PKCγ residence at the PM (13) and increase phosphorylation of PKC substrates (27, 28, 43, 49). R59022 could achieve PKC activation through the increase in DAG **(Fig. 1)** or through directly binding PKC as a lipophilic drug. Ritanserin and its putative pharmacophore fragment broadly interact with kinases, including PKCα and PKCβ (66, 67), suggesting that R59022 could bind to PKCs or a variety of other protein and lipid kinases.

PKC-mediated phosphorylation is a known driver of membrane translocation for DGKθ, and DGKζ, consistent with some of the paralog-specific recruitment we observed with R59022 treatment (18, 19, 71) **(Fig. 2)**. PKC is also known to activate the PLDs, potentially explaining the increase in PA with R59022 treatment (72, 73). Past studies have proposed that R59022 could indeed activate PLD, although others saw that a 30-minute treatment with 10 *μ*M R59022 increased the PLD substrate PC, suggesting that R59022 also inhibits PLD (27, 50). Additionally, PKC activation alone does not explain the DGKα recruitment seen, as Ca^2+^ release is also required, but we did not see any evidence of R59022 triggering Ca^2+^ release (unpublished data) (74, 75). Thus, future studies will need to determine the effect of R59022 on DAG accumulation, PKC translocation, DGK phosphorylation, and PLD activity.

Off-target interactions of R59022 with PKC or other kinases may also underlie our observed cytotoxicity in HEK293A cells. Our data is consistent with reported R59022- or ritanserin-induced apoptosis (76, 77). Although other reports note that R59022 is only toxic in cancer cell lines and not normal human astrocytes or fibroblasts (48), and two studies reported a lack of R59022 toxicity based on trypan blue assays (25, 35). Therefore, while indiscriminate kinase interactions could explain the cytotoxicity of R59022 that we see in HEK293A cells, the mechanism of this cytotoxicity and in which cell types it occurs is still unclear.

While the exact mechanism of R59022’s effects on the DGKs and PA remains unclear, we did show the effectiveness of BMS-502 in HEK293A cells **(Fig. 4)**; using BMS-502 allowed us to deconvolve DGK and PLD contributions to PA production after CCh activation of PLC **(Fig. 5)**. While BMS-502 induced PM recruitment of DGKα, DGKθ, and DGKζ; CCh only recruits DGKα to the PM **(Fig. 6)**. Thus, we suspect that it is the effects of BMS-502 against DGKα that results in the lower PA levels. Similarly, FIPI inhibits both PLD1 and PLD2, but we expect PLD2 to be the main paralog regulating PA levels as we saw constitutive PM localization regardless of CCh addition **(Fig. 6)**.

Overall, our results caution against the use of R59022 as a DGK inhibitor as it seemingly has off-target effects leading to an increase in PA levels and cytotoxicity, thus complicating interpretation of data obtained with this drug. BMS-502, however, is effective and non-toxic in HEK293A cells, as has been previously reported in Jurkat cells. Thus BMS-502 allowed us to gain important insights into the regulation of PA levels by DGKα during PLC activation.

Finally, this study further describes assays to characterize DGK and PLD activity in live cells. While the approach still has some caveats, it opens up a new avenue to study these complicated paralogs in their native environment. We anticipate future studies using this approach to look at paralog-specific roles under other stimulations or within disease models. Furthermore, we predict that these assays could be translated to other druggable lipid kinases, thus providing lipid enzymologists with a powerful new tool.

## Materials and Methods

### Cell Culture

All cell lines were grown at 37 °C with 5% CO_2_. HEK293A cells (ThermoFisher R70507) served as the parental line for HEK293A-NG2^1-^10 cells, which express strands 1-10 of NG, and CRISPR-edited HEK293A cells. Cells were maintained by being passaged twice a week. To passage, cells were washed with 1x PBS (Fisher Scientific BP3994), treated with 1 mL TrypLE (ThermoFisher 12604039), then resuspended in 4 mL complete Dulbecco’s modified Eagle’s media (DMEM: ThermoFisher 10567022, with 10% HI-FBS (ThermoFisher A5669801), 1% 100 u/mL penicillin and 100 *μ*g/mL streptomycin (ThermoFisher 15140122), and 0.1% chemically defined lipid concentrate (CDLC; ThermoFisher 11905031) before being passaged 1:5. To image the cells, 35 mm glass bottom dishes with a 20 mm aperture (CellVis D35-20-1.5-N) were coated with a mixture of 10 *μ*L 1 mg/mL ECL cell attachment matrix (Sigma 08-110) in 500 *μ*L of incomplete DMEM. Dishes were left to coat for at least 30 minutes at 37 °C. After the incubation, 2 mL warmed complete DMEM was used to replace the coating. Cells were then seeded into the dishes; enough cells were added to obtain a 100% confluent culture on the day of confocal imaging, or a 50% confluent culture on the day of TIRF imaging. After seeding, the cells were incubated for at least 2 hours at 37 °C before being transfected.

### Transfection

To overexpress proteins, 1 *μ*g of DNA was added to 100 *μ*L Opti-MEM (ThermoFisher 51985091). A solution of 3 *μ*L Lipofectamine 2000 (ThermoFisher 11668019) diluted in 100 *μ*L Opti-MEM was then added to the DNA solution, and this solution was incubated at room temperature for 5-20 minutes to form DNA/Lipofectamine complexes. After complexing, the solution was then added to each dish of cells and incubated for 4 hours at 37 °C. After that period, the transfection solution was replaced with 1.6 to 2 mL Complete HEPES-buffered Imaging Media [CHIM; 2.5% 1 M Na-HEPES buffer (VWR EM-5320) at pH 7.4, 0.1% CDLC, 1% Glutamax (ThermoFisher 35050061), and 10% HI-FBS in 500 mL Fluorobrite media (ThermoFisher A1896702)]. Once in the CHIM, the cells were then incubated at 37 °C and 5% CO_2_ until imaging the next day.

For Figure 1 and Figure 5, CellBrite® Steady 405 (Biotium 30105) or CellBrite® Steady 650 (Biotium 30108) were used to dye the plasma membrane to make a mask for confocal imaging. For these experiments, instead of replacing the transfection solution with CHIM, 1.6-2 mL of the CellBrite® dye and enhancer (both diluted 1:2000 in CHIM) was added and left on the cells for imaging the next day.

### Confocal Microscopy

A Nikon A1R confocal microscope with a Nikon TiE inverted stand was used for imaging. A stage incubator (Tokai Hit) was used to maintain the cells at 37 °C while imaging. A 100x plan apochromatic oil immersion objective was used with a 1.45 NA. Cells were excited with a dual fiber-coupled LU-NV laser launch. The channel series setting was used so 488 nm and 640 nm lasers were activated during one line scan, while a second pass activated the 561 nm and 405 nm lasers, to avoid crosstalk. Emission was collected by individual fibers: 500-550 nm (green), 663-737 nm (far-red), 570-620 nm (red), and 425-475 nm (blue).

For the pinhole, 1.2x the Airy disc size of the longest wavelength channel was used to calculate the necessary opening. To decrease the noise, some experiments utilized 8x frame averages and the Nikon Elements denoising software.

To image the cells, the positions of 10-15 fields with appropriately fluorescent cells were marked. After 2 minutes, the cells were stimulated with 500 *μ*L of the particular stimulation as indicated in **Table 1**. The concentrations listed in **Table 1** were made up as a 5x solution in CHIM, since the solutions would be diluted when added to the CHIM on the cells in the dish. When dose responses were performed, serial dilutions were used to make up 5x solutions of each dose (indicated in the figure legends), starting with a 5x solution of highest dose needed.

**Table 1.**
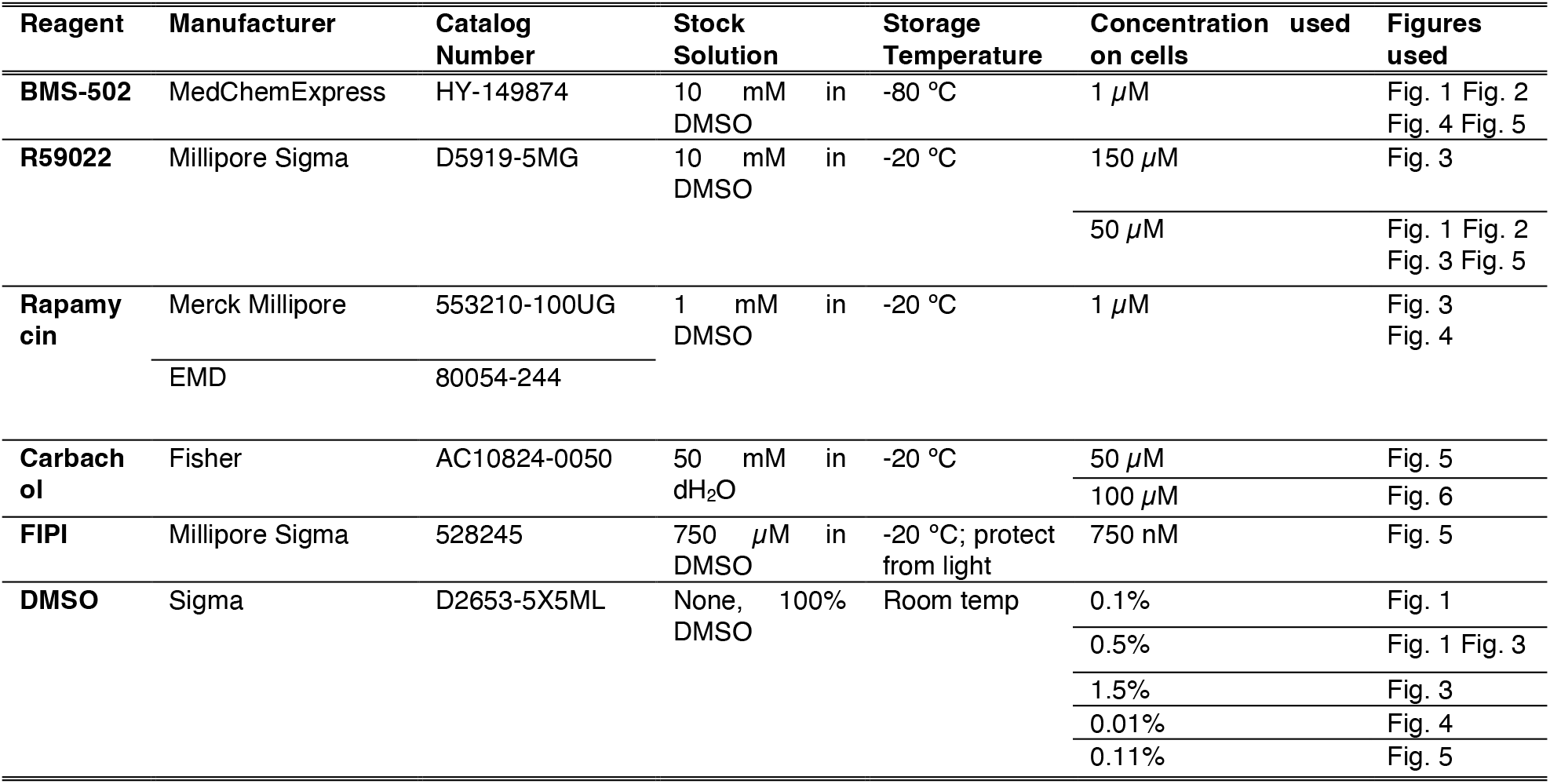
Cell stimulations used throughout this study. For the dose response curves in Figure 3 and Figure 4, the highest dose that was used in the serial dilutions is indicated in this table.

| Reagent | Manufacturer | Catalog Number | Stock Solution | Storage Temperature | Concentration used on cells | Figures used |
|---|---|---|---|---|---|---|
| **BMS-502** | MedChemExpress | HY-149874 | 10 mM in DMSO | -80 °C | 1 *µ*M | Fig. 1 Fig. 2 Fig. 4 Fig. 5 |
| **R59022** | Millipore Sigma | D5919-5MG | 10 mM in DMSO | -20 °C | 150 *µ*M | Fig. 3 |
|  |  |  |  |  | 50 *µ*M | Fig. 1 Fig. 2 Fig. 3 Fig. 5 |
| **Rapamycin** | Merck Millipore | 553210-100UG | 1 mM in DMSO | -20 °C | 1 *µ*M | Fig. 3<br>Fig. 4 |
|  | EMD | 80054-244 |  |  |  |  |
| **Carbachol** | Fisher | AC10824-0050 | 50 mM in dH<sub>2</sub>O | -20 °C | 50 *µ*M | Fig. 5 |
|  |  |  |  |  | 100 *µ*M | Fig. 6 |
| **FIPI** | Millipore Sigma | 528245 | 750 *µ*M in DMSO | -20 °C; protect from light | 750 nM | Fig. 5 |
| **DMSO** | Sigma | D2653-5X5ML | None, 100% DMSO | Room temp | 0.1% | Fig. 1 |
|  |  |  |  |  | 0.5% | Fig. 1 Fig. 3 |
|  |  |  |  |  | 1.5% | Fig. 3 |
|  |  |  |  |  | 0.01% | Fig. 4 |
|  |  |  |  |  | 0.11% | Fig. 5 |

When consecutive stimulations were used, the transfection solution was replaced with 1.6 mL CHIM. Then the first stimulation was added as a 400 *μ*L solution, and the second was added as a 500 *μ*L solution, which also included the first stimulant so as to not dilute the drug already on the cells.

For the images in **Fig. 1B**, a large static image of the field of cells was taken. To do this, 500 *μ*L of the stimulation was gently added to the cells and mixed by rocking the dish. For the 1 hour incubation, the cells were left in the 37 °C stage incubator on the confocal microscope. After the incubation period, a 1 mm x 1 mm area of the dish was imaged.

### Confocal Image Analysis

FIJI (80) was used to quantify fluorescence intensity ratios. The given ratios are either the fluorescence intensity at the PM to that within the cytosol or the fluorescence intensity at the mitochondria to that within the whole cell. In each experiment, a membrane marker was expressed, either CellBrite® for the PM or iRFP-FRB-Fis1 for the mitochondria so that the membrane could be turned into a mask where biosensor intensity could be measured to obtain the numerator in the ratio. For the denominator, a region of interest (ROI) was drawn around the whole cell or within the cytosol, while excluding any PM or nuclear fluorescence.

To create a binary mask out of the membrane marker, marker fluorescence was blurred with Gaussian filters at 1x, 2x, 3x or 4x the fluorophore’s Airy disc size. Then, the filtered images were subtracted from each other so that each image was subtracted from the next smaller filtered image. This process created wavelets which were then multiplied together, and a threshold of 0.5x the original image’s standard deviation was applied. The resulting mask was then dilated with a 1 or 2-pixel cycle and a value of 0.75x the mode of the fluorescence intensity was subtracted as background fluorescence (81). The intensity of constructs with the mask and ROI was then measured and used to produce the given ratios. Cells that were dead, dying or that moved too much during imaging were excluded from analysis. GraphPad 9 or later was used for data presentation and statistical analysis.

### Lipidomics

Cells were passaged as described above and 10,000,000 cells were seeded into 10 mL warmed complete DMEM in a 10 cm^2^ dish. The dish was incubated at 37 °C overnight. The next day, the media was aspirated, and the cells were washed 3x with 10 mL CDLC-free serum-free CHIM (1% Glutamax and 2.5% 1 M Na-HEPES-buffer in Fluorobrite media). The final wash was replaced with either 0.5% DMSO or 50 *μ*M R59022 in 5 mL CDLC-free serum-free CHIM. Cells were treated at their respective conditions for 5 minutes at 37 °C. The plates were then transferred to ice and cells were scraped off and centrifuged for 3 min at 500 x g. The pellet was resuspended in 1 mL ice cold 1x PBS and transferred to a cryo-vial. Cells were centrifuged again for 3 min at 500 x g, and the PBS was aspirated until the pellet was fully dried. Cells were then flash frozen in liquid nitrogen and stored at −80 °C or on dry ice.

Lipid extraction was performed as previously described, with modifications (57). Briefly, lipids were extracted from 10 × 10^6^ HEK293 cells using a Bligh–Dyer protocol (CHCl_3_/MeOH/H_2_O, 1:2:1). Cell pellets were resuspended in H_2_O (1 mL) and added to CHCl_3_ (1 mL) and MeOH (2 mL) containing butylated hydroxytoluene (50 *μ*g mL^−1^). Deuterated internal standards (15 *μ*L 1:1 LipidSplash Lipidomix per sample) were added to organic solvents prior to extraction. Samples were vortexed, incubated on ice (20 min), and centrifuged (2,000 × g, 5 min, 4 °C) to separate phases. The organic phase was collected, and the aqueous phase was re-extracted using an acidified modified Folch protocol (CHCl_3_/MeOH/H_2_O, 2:1:1, final 1 M HCl). Combined organic extracts were dried under nitrogen and resuspended in methanol:isopropanol (1:1, v/v) for storage at −80 °C prior to analysis.

Low-microflow targeted lipidomics was performed as previously described (58). ESI–LC–MS/MS was performed on a TSQ Quantis Plus triple quadrupole mass spectrometer coupled to a Vanquish Neo UHPLC system equipped with a C12 column (0.5 mm i.d. × 150 mm) maintained at 50 °C. One microliter of sample was injected. Mobile phases consisted of A (50% ACN, 50% H_2_O, 0.1% formic acid, 10 mM ammonium formate) and B (88% IPA, 10% ACN, 2% H_2_O, 0.1% formic acid, 10 mM ammonium formate), delivered at 25 *μ*L min^−1^. The instrument was operated in positive and negative ion mode to detect DAGs and PAs, respectively.

### CRISPR/Cas9 Knock-in Cell Line Generation

Designs for CRISPR HDRTs and gRNA are shown in **Table 2**. For PLD2 tagging, we used the HDRT, gRNA and genotyping primers already utilized in the OpenCell database. For PLD1, we used protoSpaceJAM to design the HDRTs, gRNAs and genotyping primers (60, 82). HDRTs given at high concentrations were reconstituted in nuclease-free water to 100 pmol/*μ*L. Low concentration HDRTs were not resuspended before using. gRNAs were re-constituted to 20 pmol/*μ*L in nuclease-free water. HDRTs and gRNAs were stored at −20 °C until use.

**Table 2.**
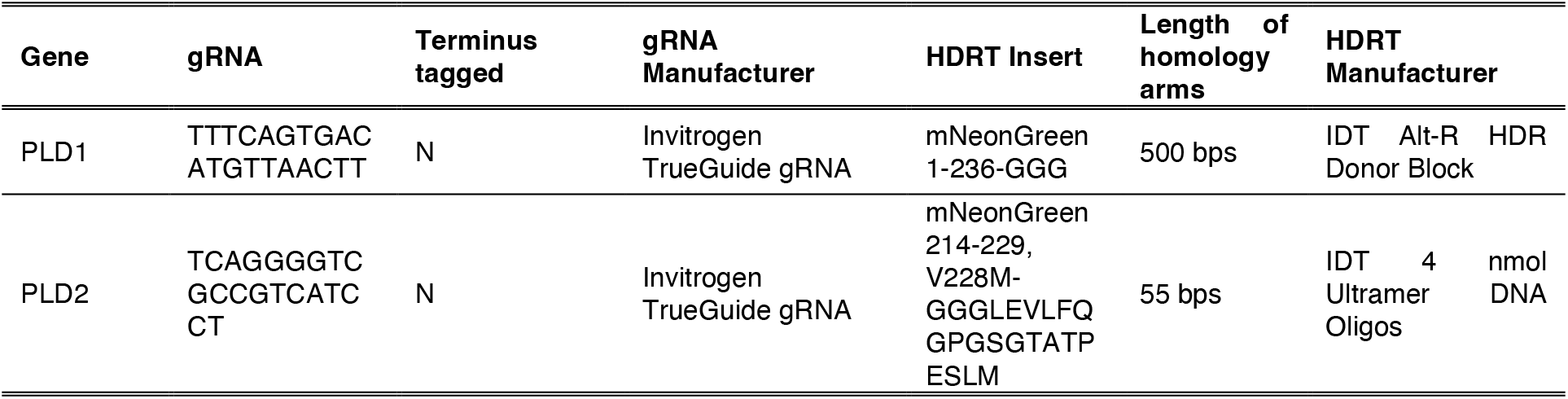
gRNA and HDRT sequences for insertion of fluorophores by CRISPR/Cas9. gRNA sequences are shown by nucleotides, while HDRT sequences are shown by amino acids. Numbering of amino acids follows that of the full-length protein. The cut site for N-terminal tagging was immediately after the start codon.

Electroporation of gRNA and HDRTs was done using the Neon Transfection System (ThermoFisher MPK1096). The gRNA complex was made up of 10 pmol gRNA, 10 pmol TrueCUT Cas9 protein V2 (ThermoFisher A36498), and 5 *μ*L Buffer R. This mixture was incubated for at least 5 mins at room temperature. HEK293A cells or HEK293A-NG2^1-10^ cells were then passaged and pelleted as described in the Neon Transfection System protocol. The cells were resuspended in Buffer R to create a solution of 40,000 cells/*μ*L.

For HDRTs that were resuspended, 100 pmol of the HDRT was added to the gRNA complex. For HDRTs that remained as powder, the tube was spun briefly. Then, the gRNA complex was added to the HDRT tube and mixed by gentle pipetting.

To electroporate the cells, 5 *μ*L of the cell suspension was added to the gRNA and HDRT complex and mixed gently. 10 *μ*L of this solution was then aspirated into a Neon tip and electroporated at 1500 V for 1 pulse at 20 ms. The electroporated cells were then added to a pre-warmed 6-well plate containing 2 mL antibiotic-free complete DMEM (complete DMEM without penicillin or streptomycin). In some experiments, 1.7 *μ*L of Alt-R^TM^ HDR Enhancer V2 (IDT 10007921) was included in the wells. Cells were then incubated at 37 °C for 12-16 hours before the antibiotic-free DMEM was replaced with complete DMEM.

Once the wells were fully confluent, each well was passaged, and cells were seeded into pre-coated glass bottom dishes as described above. The rest of the cells not seeded into dishes were passaged into a new flask to continue to grow. A 1 mm x 1 mm field of the cells in the dish was imaged on the confocal microscope using 15% 488 nm (green) laser power with 35 gain and 0% 561 nm (red) laser power with 35 gain. The channel series setting was turned off in order to capture any autofluorescence. 16x line averaging was used to help reduce noise. After imaging, the green image was divided by the red image using FIJI and the resulting ratio image was used to quantify the number of cells showing fluorescence in the green channel, as these were considered to be the tagged cells. The total number of cells was counted using the DIC image and these counts were used to calculate the percent efficiency of tag integration.

To sort the flask of the mixed population of electroporated cells, the cells were passaged, centrifuged for 3 min at 500 x g, and resuspended in 200 *μ*L CHIM using a sterile 40 *μ*m nylon mesh cell strainer (Fisherbrand 22363547). The cells were then sorted using FACS (University of Pittsburgh Department of Immunology Unified FLOW Core). The percent efficiency as calculated by confocal imaging was used to gate the sorting. Sorted cells were seeded into a 6-well plate with 2 mL pre-warmed complete DMEM. The process of confocal screening and FLOW sorting was repeated as needed in order to get a population where 95-100% of cells were tagged.

### Genotyping PCR

Genomic DNA was extracted from tagged and parental cell lines using the PureLink Genomic DNA Mini Kit (ThermoFisher K182001). DNA was diluted to 10 ng/*μ*L in dH_2_O and stored as aliquots at −20 °C. Primers used for genotyping are described in **Table 3**. All primers were synthesized by Thermo Fisher and stored as 100 *μ*M solutions. The PCR mix was made up using 2.5 *μ*L forward primer, 2.5 *μ*L reverse primer, 20 *μ*L genomic DNA, and 25 *μ*L 2x DreamTaq Green PCR Master Mix (Thermo Scientific K1081). PCR was run using the following steps: 95 °C for 2 min, 95 °C for 30 s, annealing temperature for 30 s (see **Table 3**), 72 °C for 60 s, go back to step 2 x 34, 72 °C for 10 min, 4 °C hold. PCR products were then run on a 2% agarose gel for 30-60 mins at 95 V. The expected band was then excised and purified using the GeneJet Gel Extraction and DNA Cleanup Kit (Fisher Scientific FERK0832).

**Table 3.**
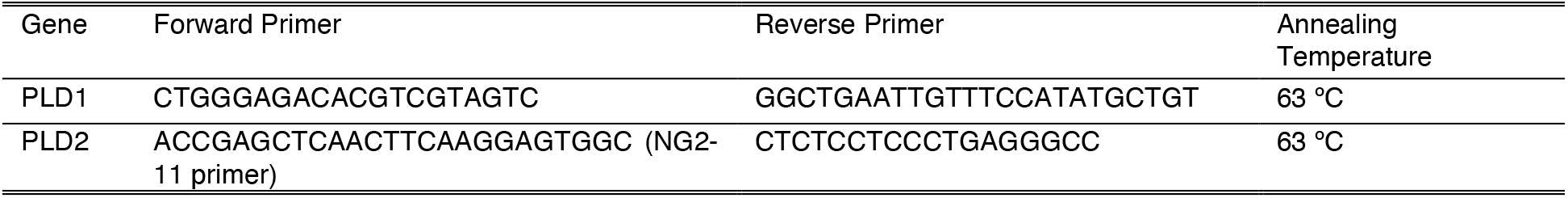
Genotyping PCR Conditions. The following forward and reverse primers were used in PCR with the respective genomic DNA and the annealing temperature listed. These primers were also used to sequence the PCR products.

The purified band was then re-amplified to increase the concentration. The second PCR mix utilized 1 *μ*L forward primer, 1 *μ*L reverse primer, 8 *μ*L extracted DNA and 10 *μ*L 2x DreamTaq. The same reaction conditions were used as described above, except in some cases the reaction was increased to 44 cycles. After PCR, the product was column purified using the GeneJet Gel Extraction and DNA Cleanup Kit. This additional PCR and purification process was repeated until a concentration of 2-10 ng/*μ*L was reached. 20 ng product in 10 *μ*L dH_2_O with 25 pmol of each genotyping primer was then sent for Sanger sequencing (GENEWIZ from Azenta Life Sciences) to confirm integration of the fluorophore.

### TIRF Microscopy

A Nikon TiE inverted microscope with Nikon TIRF illuminator was used for imaging. The microscope included a chamber heated to 37 °C to incubate cells during imaging experiments. Cells were excited using an Oxxius L4C laser launch, and imaged with a 100× 1.45 NA plan apochromatic oil-immersion objective. Fields of 1024 × 1024 pixels, using 2× pixel binning, were imaged with a 1.5× magnifier. To collect images, a Hamamatsu ORCA-Fusion BT sCMOS camera was used with ultra-low noise mode and 2× pixel binning.

While imaging, 2-5 fields were selected with cells that showed optimized protein expression. NeonGreen and StayGold fluorophores on endogenous proteins were excited with the 488 nm laser at 30% power and 100 ms exposure. The resulting fluorescence emission passed through a 500-550 nm filter. Cells were imaged every 5 s for the duration of the experiment, which consisted of 2 minutes of baseline imaging and then several minutes of imaging after the addition of 500 *μ*L of the appropriate stimulant as indicated in **Table 1**.

### TIRF Image Analysis

To quantify single particles of proteins associated with the PM, a 10 *μ*m x 10 *μ*m square ROI was drawn in a representative area of the cell using FIJI (80). The ThunderSTORM plugin (83) was then used to count the fluorescent particles within this ROI at each time point. To visualize the single particles in the images presented in this manuscript, a rolling-ball average of 10 pixels was used to subtract the background. Graphs and statistics were produced using GraphPad 9 or later.

### Plasmids and Cloning

Plasmids used throughout this study are shown in **Table 4**. Plasmids were made through either PCR amplification of an insert followed by NEB HiFi Gibson Assembly (New England Biolabs E5520S) or by restriction digest (FastDigest Thermo Scientific) and ligation (T4 DNA Ligase Thermo Scientific EL0011). Insert sequences were obtained through previously generated plasmids or by a custom GeneBlock (IDT). All plasmid sequences were confirmed by Sanger sequencing over the inserted sequence or with long-read nanopore sequencing over the full plasmid (GENEWIZ from Azenta Life Sciences).

**Table 4.** Plasmids used in this study. All genes are *Homo sapiens* unless indicated otherwise. The relevant amino acid positions in the full-length protein are noted. Mutations are described by the position of the residues in the full-length protein. Amino acid sequences written out indicate linkers.

| Name | Sequence | Reference |
|---|---|---|
| mCh-C1ab-Prkd1 | *X.leavis* map2k1.L<sup>32-44</sup>-APVAT-mCherry-SGLRSRAQASNSTS-*M. musculus* Prkd1<sup>138-343, T208A, N313D</sup> | (84) |
| NG-PILS-Nir1 | NeonGreen-GGSGGM-PITPNM3 paralog 1<sup>613-897, A885V</sup> | (4) |
| iRFP-FRB-Fis1 | iRFP-SGLRSRAGGAGAILSR-MTOR<sup>2021-2113</sup>-GGSAGGSAQASNSAVDGTA-Fis1<sup>122-152</sup> | (85) |
| mRFP-FKBP-DGKa-cat | mRFP-SGLRSRSAAAGAGGAARAAL-FKBP1A<sup>3-108</sup>-SAGGSAGGSAGGSAGGSAGGPRAQASRS-DGKA paralog a<sup>356-735</sup> | (4) |
| TagBFP2-FKBPflex-*Bc*-PI-PLC | *B. cereus* PI-PLC<sup>32-329, W78A, W273A</sup>-RILQSTVPMG-FKBP1A<sup>3-108</sup>-RDPPVATM-TagBFP2-SGLRSRSAAATLDHNQPYHICRGFTCFKKPPTPPPEPET | (86) |
| iRFP-SEC61B | iRFP713-SGLRS-SEC61B<sup>1-96</sup> | (87) |
| pUC19 | Empty plasmid used to bring total DNA mass up to 1 µg as needed | (88) |

## Data availability

All data generated or analyzed during this study are included in this published article and its supplementary information files.

## Author Contributions

Conceptualization; Visualization; Writing - Original Draft; Writing - Review & Editing: Claire C. Weckerly. Data analysis; Investigation: Visualization; Writing – Review & Editing: Tiernan Swayhoover, Olivia Murtagh. Methodology: Joshua G. Pemberton. Conceptualization; Data analysis; Writing - Review & Editing: Ku-Lung Hsu, Gerald R.V. Hammond.

K.-L.H. is a founder and scientific advisory board member of Hyku Biosciences. The other authors declare no competing financial interests.

---

**Supplemental Figure 1.**
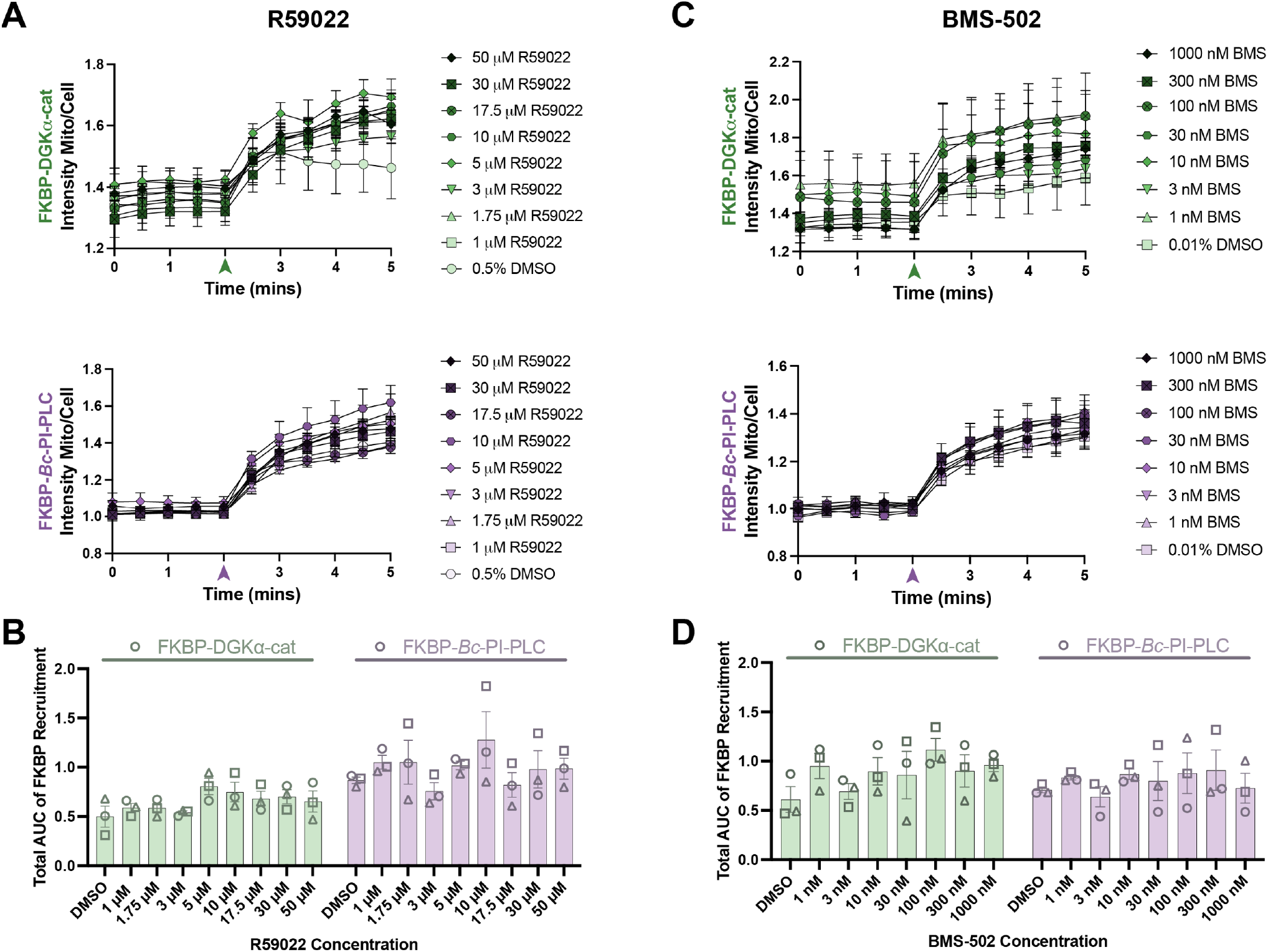
DGK inhibitors do not affect recruitment of FKBP constructs to the mitochondria. **(A)** Recruitment of FKBP-DGKacat and FKBP-*Bc*-PI-PLC to the mitochondria after the addition of rapamycin and increasing doses of R59022 at the timepoint indicated by the arrow. **(B)** Total area under the curve of the recruitment shown in panel A. **(C)** As in panel A, but with increasing doses of BMS-502. **(D)** As in panel B, but showing the area under the curve data from panel C. Data in panel A-B is taken from the same cells as in Fig. 3C and data in panel C-D is from the cells in Fig. 4B. Timelapse data shown is the grand means of 3 independent experiments ± SEM. Bar graphs show the mean ± SEM of 3 independent experiments. Data from the same replicates are matched by symbol shape. 30-44 cells were analyzed.

## References

1. Cockcroft, S., and Raghu, P. (2016) Topological organisation of the phosphatidylinositol 4,5-bisphosphate–phospholipase C resynthesis cycle: PITPs bridge the ER–PM gap. Biochem J. 473, 4289–4310

2. Nobe, K., Ohata, H., and Momose, K. (1994) Activation of diacylglycerol kinase by carbachol in guinea pig taenia coli. Biochem. Pharmacol. 48, 2005–2014

3. Liang, D., Wu, K., Tei, R., Bumpus, T. W., Ye, J., and Baskin, J. M. (2019) A real-time, click chemistry imaging approach reveals stimulus-specific subcellular locations of phospholipase D activity. Proc. Natl. Acad. Sci. 116, 15453–15462

4. Weckerly, C. C., Rahn, T. A., Ehrlich, M., Wills, R. C., Pemberton, J. G., Airola, M. V., and Hammond, G. R. V. (2025) PILS-Nir1 is a sensitive phosphatidic acid biosensor that reveals mechanisms of lipid production. J. Cell Biol. 224, e202405174

5. Touyz, R. M., Alves-Lopes, R., Rios, F. J., Camargo, L. L., Anagnostopoulou, A., Arner, A., and Montezano, A. C. (2018) Vascular smooth muscle contraction in hypertension. Cardiovasc Res. 114, 529–539

6. Peng, Z., and Arendshorst, W. J. (2007) Phospholipase C Expression and Activity in Smooth Muscle Cells of Renal Arterioles and Aorta of Young, Spontaneously Hypertensive Rats During Culture. Am J Hypertens. 20, 520–526

7. Peng, Z., Dang, A., and Arendshorst, W. J. (2007) Increased Expression and Activity of Phospholipase C in Renal Arterioles of Young Spontaneously Hypertensive Rats. Am J Hypertens. 20, 38–43

8. Sakane, F., Hoshino, F., Ebina, M., Sakai, H., and Takahashi, D. (2021) The Roles of Diacylglycerol Kinase α in Cancer Cell Proliferation and Apoptosis. Cancers. 13, 5190

9. Sakane, F., Imai, S., Kai, M., Yasuda, S., and Kanoh, H. (2008) Diacylglycerol Kinases as Emerging Potential Drug Targets for a Variety of Diseases. Curr Drug Targets. 9, 626–640

10. Sakane, F., Mizuno, S., and Komenoi, S. (2016) Diacylglycerol Kinases as Emerging Potential Drug Targets for a Variety of Diseases: An Update. Frontiers Cell Dev Biology. 4, 82

11. Sakane, F., Imai, S., Kai, M., Yasuda, S., and Kanoh, H. (2007) Diacylglycerol kinases: Why so many of them? Biochimica Et Biophysica Acta Bba - Mol Cell Biology Lipids. 1771, 793–806

12. Selvy, P. E., Lavieri, R. R., Lindsley, C. W., and Brown, H. A. (2011) Phospholipase D: Enzymology, Functionality, and Chemical Modulation. Chem Rev. 111, 6064–6119

13. Shirai, Y., Segawa, S., Kuriyama, M., Sakai, N., Saito, N., and Goto, K. (2000) Subtype-specific Translocation of Diacylglycerol Kinase α and γ and Its Correlation with Protein Kinase C*. J. Biol. Chem. 275, 24760–24766

14. Sanjuán, M. A., Pradet-Balade, B., Jones, D. R., Martínez-A, C., Stone, J. C., Garcia-Sanz, J. A., and Mérida, I. (2003) T Cell Activation In Vivo Targets Diacylglycerol Kinase α to the Membrane: A Novel Mechanism for Ras Attenuation. J. Immunol. 170, 2877–2883

15. Sanjuán, M. A., Jones, D. R., Izquierdo, M., and Mérida, I. (2001) Role of Diacylglycerol Kinase α in the Attenuation of Receptor Signaling. J Cell Biology. 153, 207–220

16. Imai, S., Sakane, F., and Kanoh, H. (2002) Phorbol Ester-regulated Oligomerization of Diacylglycerol Kinase δ Linked to Its Phosphorylation and Translocation*. J. Biol. Chem. 277, 35323–35332

17. Imai, S., Kai, M., Yamada, K., Kanoh, H., and Sakane, F. (2004) The plasma membrane translocation of diacylglycerol kinase δ1 is negatively regulated by conventional protein kinase C-dependent phosphorylation at Ser-22 and Ser-26 within the pleckstrin homology domain. Biochem J. 382, 957–966

18. Santos, T., Carrasco, S., Jones, D. R., Mérida, I., and Eguinoa, A. (2002) Dynamics of Diacylglycerol Kinase ζ Translocation in Living T-cells STUDY OF THE STRUCTURAL DOMAIN REQUIREMENTS FOR TRANSLOCATION AND ACTIVITY* 210. J Biol Chem. 277, 30300–30309

19. Baal, J. van, Widt, J. de, Divecha, N., and Blitterswijk, W. J. van (2005) Translocation of Diacylglycerol Kinase θ from Cytosol to Plasma Membrane in Response to Activation of G Protein-coupled Receptors and Protein Kinase C*. J Biol Chem. 280, 9870–9878

20. Brown, F. D., Thompson, N., Saqib, K. M., Clark, J. M., Powner, D., Thompson, N. T., Solari, R., and Wakelam, M. J. O. (1998) Phospholipase D1 localises to secretory granules and lysosomes and is plasma-membrane translocated on cellular stimulation. Curr. Biol. 8, 835–838

21. Du, G., Altshuller, Y. M., Vitale, N., Huang, P., Chasserot-Golaz, S., Morris, A. J., Bader, M.-F., and Frohman, M. A. (2003) Regulation of phospholipase D1 subcellular cycling through coordination of multiple membrane association motifs. J Cell Biology. 162, 305–315

22. Freyberg, Z., Sweeney, D., Siddhanta, A., Bourgoin, S., Frohman, M., and Shields, D. (2001) Intracellular Localization of Phospholipase D1 in Mammalian Cells. Mol. Biol. Cell. 12, 943–955

23. Colley, W. C., Sung, T.-C., Roll, R., Jenco, J., Hammond, S. M., Altshuller, Y., Bar-Sagi, D., Morris, A. J., and Frohman, M. A. (1997) Phospholipase D2, a distinct phospholipase D isoform with novel regulatory properties that provokes cytoskeletal reorganization. Curr. Biol. 7, 191–201

24. Sciorra, V. A., Rudge, S. A., Wang, J., McLaughlin, S., Engebrecht, J., and Morris, A. J. (2002) Dual role for phosphoinositides in regulation of yeast and mammalian phospholipase D enzymes. J. Cell Biol. 159, 1039–1049

25. Flores, I., Casaseca, T., Martinez-A, C., Kanoh, H., and Merida, I. (1996) Phosphatidic Acid Generation through Interleukin 2 (IL-2)-induced α-Diacylglycerol Kinase Activation Is an Essential Step in IL-2-mediated Lymphocyte Proliferation (∗). J. Biol. Chem. 271, 10334–10340

26. Courcelles, D. de C. de, Roevens, P., Belle, H. V., Kennis, L., Somers, Y., and Clerck, F. D. (1989) The Role of Endogenously Formed Diacylglycerol in the Propagation and Termination of Platelet Activation A biochemical and functional analysis using the novel diacylglycerol kinase inhibitor, R 59 949. J Biol Chem. 264, 3274–3285

27. Courcelles, D. C. de C. de, Roevens, P., and Belle, H. V. (1985) R 59 022, a diacylglycerol kinase inhibitor. Its effect on diacylglycerol and thrombin-induced C kinase activation in the intact platelet. J Biological Chem. 260, 15762–70

28. Boroda, S., Niccum, M., Raje, V., Purow, B. W., and Harris, T. E. (2017) Dual activities of ritanserin and R59022 as DGKα inhibitors and serotonin receptor antagonists. Biochem. Pharmacol. 123, 29–39

29. Jiang, Y., Sakane, F., Kanoh, H., and Walsh, J. P. (2000) Selectivity of the diacylglycerol kinase inhibitor 3-{2-(4-[bis-(4-fluorophenyl)methylene]-1-piperidinyl)ethyl}-2,3-dihydro-2-thioxo-4(1H)quinazolinone (R59949) among diacylglycerol kinase subtypes. Biochem Pharmacol. 59, 763–772

30. Sakane, F., Yamada, K., and Kanoh, H. (1989) Different effects of sphingosine, R59022 and anionic amphiphiles on two diacylglycerol kinase isozymes purified from porcine thymus cytosol. FEBS Lett. 255, 409–413

31. Sato, M., Liu, K., Sasaki, S., Kunii, N., Sakai, H., Mizuno, H., Saga, H., and Sakane, F. (2013) Evaluations of the Selectivities of the Diacylglycerol Kinase Inhibitors R59022 and R59949 Among Diacylglycerol Kinase Isozymes Using a New Non-Radioactive Assay Method. Pharmacology. 92, 99–107

32. Ascher-Landsberg, J., Saunders, T., and Phillippe, M. (2000) The role of diacylglycerol as a modulator of oxytocin-stimulated phasic contractions in myometrium from pregnant and nonpregnant rats. Am. J. Obstet. Gynecol. 182, 943–949

33. Aragonés, J., Jones, D. R., Martiń, S., Juan, M. A. S., Alfranca, A., Vidal, F., Vara, A., Mérida, I., and Landázuri, M. O. (2001) Evidence for the Involvement of Diacylglycerol Kinase in the Activation of Hypoxia-inducible Transcription Factor 1 by Low Oxygen Tension*. J. Biol. Chem. 276, 10548–10555

34. Dale, M. M., and Penfield, A. (1987) Comparison of the effects of indomethacin, RHC80267 and R59022 on superoxide production by 1,oleoyl-2,acetyl glycerol and A23187 in human neutrophils. Br. J. Pharmacol. 92, 63–68

35. Ederveen, A. G. H., Vries, S. E. van E., Pont, J. J. H. H. M. D., and Willems, P. H. G. M. (1990) The diacylglycerol kinase inhibitor, R59022, potentiates cholecystokinin-induced enzyme secretion from rabbit pancreatic acini. Eur. J. Biochem. 188, 333–338

36. Ford, C. P., Stemkowski, P. L., Light, P. E., and Smith, P. A. (2003) Experiments to Test the Role of Phosphatidylinositol 4,5-Bisphosphate in Neurotransmitter-Induced M-Channel Closure in Bullfrog Sympathetic Neurons. J. Neurosci. 23, 4931–4941

37. Gomez-Cambronero, J., Molski, T. F. P., Becker, E. L., and Sha’afi, R. I. (1987) The diacylglycerol kinase inhibitor R59022 potentiates superoxide production but not secretion induced by fMet-Leu-Phe: Effects of leupeptin and the protein kinase C inhibitor H-7. Biochem. Biophys. Res. Commun. 148, 38–46

38. Hurttia, H., and Leino, L. (1996) Subcellular localization of diacylglycerol kinase activity in stimulated and unstimulated human peripheral blood lymphocytes and neutrophils. IUBMB Life. 40, 579–585

39. Lingelem, A. B. D., Kavaliauskiene, S., Halsne, R., Klokk, T. I., Surma, M. A., Klose, C., Skotland, T., and Sandvig, K. (2021) Diacylglycerol kinase and phospholipase D inhibitors alter the cellular lipidome and endosomal sorting towards the Golgi apparatus. Cell Mol Life Sci. 78, 985–1009

40. Mege, J. L., Tao, W., Molski, T. F., Gomez-Cambronero, J., Huang, C. K., Becker, E. L., and Sha’afi, R. I. (1988) Diacylglycerol kinase inhibitor R59022 and stimulated neutrophil responses. Am. J. Physiol.-Cell Physiol. 255, C589–C594

41. Muid, R. E., Penfield, A., and Dale, M. M. (1987) The diacylglycerol kinase inhibitor, R59022, enhances the superoxide generation from human neutrophils induced by stimulation of fMet-Leu-Phe, IgG and C3b receptors. Biochem. Biophys. Res. Commun. 143, 630–637

42. Nobe, K., Miyatake, M., Nobe, H., Sakai, Y., Takashima, J., and Momose, K. (2004) Novel diacylglycerol kinase inhibitor selectively suppressed an U46619-induced enhancement of mouse portal vein contraction under high glucose conditions. Br. J. Pharmacol. 143, 166–178

43. Ohtsuka, T., Hiura, M., Yoshida, K., Okamura, N., and Ishibashi, S. (1990) A diacylglycerol kinase inhibitor, R 59 022, potentiates superoxide anion production and 46-kDa protein phosphorylation in guinea pig polymorphonuclear leukocytes. J. Biol. Chem. 265, 15418–23

44. Orlati, S., Porcelli, A. M., Hrelia, S., Lorenzini, A., and Rugolo, M. (1998) Intracellular calcium mobilization and phospholipid degradation in sphingosylphosphorylcholine-stimulated human airway epithelial cells. Biochem. J. 334, 641–649

45. Rider, L. G., and Niedel, J. E. (1987) Diacylglycerol accumulation and superoxide anion production in stimulated human neutrophils. J. Biol. Chem. 262, 5603–8

46. Ruffo, E., Malacarne, V., Larsen, S. E., Das, R., Patrussi, L., Wülfing, C., Biskup, C., Kapnick, S. M., Verbist, K., Tedrick, P., Schwartzberg, P. L., Baldari, C. T., Rubio, I., Nichols, K. E., Snow, A. L., Baldanzi, G., and Graziani, A. (2016) Inhibition of diacylglycerol kinase α restores restimulation-induced cell death and reduces immunopathology in XLP-1. Sci. Transl. Med. 8, 321ra7

47. Burt, R. P., Chappie, C. R., and Marshall, I. (1996) The role of diacylglycerol and activation of protein kinase C in α1A-adrenoceptor-mediated contraction to noradrenaline of rat isolated epididymal vas deferens. Br. J. Pharmacol. 117, 224–230

48. Dominguez, C. L., Floyd, D. H., Xiao, A., Mullins, G. R., Kefas, B. A., Xin, W., Yacur, M. N., Abounader, R., Lee, J. K., Wilson, G. M., Harris, T. E., and Purow, B. W. (2013) Diacylglycerol Kinase α Is a Critical Signaling Node and Novel Therapeutic Target in Glioblastoma and Other Cancers. Cancer Discov. 3, 782–797

49. Nunn, D. L., and Watson, S. P. (1987) A diacylglycerol kinase inhibitor, R59022, potentiates secretion by and aggregation of thrombin-stimulated human platelets. Biochem. J. 243, 809–813

50. Mahadevappa, V. G. (1988) [3H]Phosphatidic acid formed in response to FMLP is not inhibited by R59 022, a diacylglycerol kinase inhibitor. Biochem. Biophys. Res. Commun. 153, 1097–1104

51. Bohdanowicz, M., Schlam, D., Hermansson, M., Rizzuti, D., Fairn, G. D., Ueyama, T., Somerharju, P., Du, G., and Grinstein, S. (2013) Phosphatidic acid is required for the constitutive ruffling and macropinocytosis of phagocytes. Mol Biol Cell. 24, 1700–1712

52. Chang, C.-L., and Liou, J. (2015) Phosphatidylinositol 4,5-Bisphosphate Homeostasis Regulated by Nir2 and Nir3 Proteins at Endoplasmic Reticulum-Plasma Membrane Junctions*. J Biol Chem. 290, 14289–14301

53. Kim, Y. J., Guzman-Hernandez, M.-L., Wisniewski, E., and Balla, T. (2015) Phosphatidylinositol-Phosphatidic Acid Exchange by Nir2 at ER-PM Contact Sites Maintains Phosphoinositide Signaling Competence. Dev Cell. 33, 549–561

54. Zhang, F., Wang, Z., Lu, M., Yonekubo, Y., Liang, X., Zhang, Y., Wu, P., Zhou, Y., Grinstein, S., Hancock, J. F., and Du, G. (2014) Temporal Production of the Signaling Lipid Phosphatidic Acid by Phospholipase D2 Determines the Output of Extracellular Signal-Regulated Kinase Signaling in Cancer Cells. Mol Cell Biol. 34, 84–95

55. Chupak, L., Wichroski, M., Zheng, X., Ding, M., Martin, S., Allard, C., Shi, J., Gentles, R., Meanwell, N. A., Fang, J., Tenney, D., Tokarski, J., Cao, C., and Wee, S. (2023) Discovery of Potent, Dual-Inhibitors of Diacylglycerol Kinases Alpha and Zeta Guided by Phenotypic Optimization. ACS Med. Chem. Lett. 14, 929–935

56. Wichroski, M., Benci, J., Liu, S.-Q., Chupak, L., Fang, J., Cao, C., Wang, C., Onorato, J., Qiu, H., Shan, Y., Banas, D., Powles, R., Locke, G., Witt, A., Stromko, C., Qi, H., Zheng, X., Martin, S., Ding, M., Gentles, R., Meanwell, N., Velaparthi, U., Olson, R., Wee, S., Tenney, D., Parker, C. G., Cravatt, B. F., Lawrence, M., Borzilleri, R., and Lees, E. (2023) DGKα/ζ inhibitors combine with PD-1 checkpoint therapy to promote T cell–mediated antitumor immunity. Sci. Transl. Med. 15, eadh1892

57. Shaikh, M., Mookherjee, S. P., Weckerly, C. C., Libby, A. H., Xiao, A., Zhao, Y., Vaidya, S. D., Kim, A., Li, Z., Ware, M. L., Marants, M., Murtagh, O. L., Wolfe, W. J., Bullock, T. N., Purow, B. W., Hammond, G. R. V., and Hsu, K.-L. (2025) Stereoselective Degradation of Diacylglycerol Kinases Potentiate T cell Activation and Tumor Cell Cytotoxicity. bioRxiv. 10.64898/2025.12.09.692983

58. Murtagh, OliviaL., Lowrey, P. E., and Hsu, K.-L. (2026) Ultrasensitive Detection of Tissue Lipids at the Femtomole Scale Using Low-Microflow Targeted Lipidomics. Anal. Chem. 98, 5250–5260

59. Ware, T. B., Franks, C. E., Granade, M. E., Zhang, M., Kim, K.-B., Park, K.-S., Gahlmann, A., Harris, T. E., and Hsu, K.-L. (2020) Reprogramming fatty acyl specificity of lipid kinases via C1 domain engineering. Nat. Chem. Biol. 16, 170–178

60. Cho, N. H., Cheveralls, K. C., Brunner, A.-D., Kim, K., Michaelis, A. C., Raghavan, P., Kobayashi, H., Savy, L., Li, J. Y., Canaj, H., Kim, J. Y. S., Stewart, E. M., Gnann, C., McCarthy, F., Cabrera, J. P., Brunetti, R. M., Chhun, B. B., Dingle, G., Hein, M. Y., Huang, B., Mehta, S. B., Weissman, J. S., Gómez-Sjöberg, R., Itzhak, D. N., Royer, L. A., Mann, M., and Leonetti, M. D. (2022) OpenCell: Endogenous tagging for the cartography of human cellular organization. Science. 375, eabi6983

61. Su, W., Yeku, O., Olepu, S., Genna, A., Park, J.-S., Ren, H., Du, G., Gelb, M. H., Morris, A. J., and Frohman, M. A. (2009) 5-Fluoro-2-indolyl des-chlorohalopemide (FIPI), a Phospholipase D Pharmacological Inhibitor That Alters Cell Spreading and Inhibits Chemotaxis. Mol Pharmacol. 75, 437–446

62. Monovich, L., Mugrage, B., Quadros, E., Toscano, K., Tommasi, R., LaVoie, S., Liu, E., Du, Z., LaSala, D., Boyar, W., and Steed, P. (2007) Optimization of Halopemide for Phospholipase D2 inhibition. Bioorg Med Chem Lett. 17, 2310–2311

63. Feng, S., Sekine, S., Pessino, V., Li, H., Leonetti, M. D., and Huang, B. (2017) Improved split fluorescent proteins for endogenous protein labeling. Nat. Commun. 8, 370

64. Glukhov, E., Shulga, Y. V., Epand, R. F., Dicu, A. O., Topham, M. K., Deber, C. M., and Epand, R. M. (2007) Membrane interactions of the hydrophobic segment of diacylglycerol kinase epsilon. Biochim. Biophys. Acta (BBA) - Biomembr. 1768, 2549–2558

65. Matsui, H., Hozumi, Y., Tanaka, T., Okada, M., Nakano, T., Suzuki, Y., Iseki, K., Kakehata, S., Topham, M. K., and Goto, K. (2014) Role of the N-terminal hydrophobic residues of DGKε in targeting the endoplasmic reticulum. Biochim. Biophys. Acta (BBA) - Mol. Cell Biol. Lipids. 1841, 1440–1450

66. Franks, C. E., Campbell, S. T., Purow, B. W., Harris, T. E., and Hsu, K.-L. (2017) The Ligand Binding Landscape of Diacylglycerol Kinases. Cell Chem Biol. 24, 870–880.e5

67. Huang, T., Hosseinibarkooie, S., Borne, A. L., Granade, M. E., Brulet, J. W., Harris, T. E., Ferris, H. A., and Hsu, K.-L. (2021) Chemoproteomic profiling of kinases in live cells using electrophilic sulfonyl triazole probes. Chem. Sci. 12, 3295–3307

68. Mendez, R., Shaikh, M., Lemke, M. C., Yuan, K., Libby, A. H., Bai, D. L., Ross, M. M., Harris, T. E., and Hsu, K.-L. (2023) Predicting small molecule binding pockets on diacylglycerol kinases using chemoproteomics and AlphaFold. RSC Chem. Biol. 4, 422–430

69. Das, J., and Rahman, G. M. (2014) C1 Domains: Structure and Ligand-Binding Properties. Chem. Rev. 114, 12108–12131

70. Hurley, J. H., Newton, A. C., Parker, P. J., Blumberg, P. M., and Nishizuka, Y. (1997) Taxonomy and function of C1 protein kinase C homology domains. Protein Sci. 6, 477–480

71. Abramovici, H., Hogan, A. B., Obagi, C., Topham, M. K., and Gee, S. H. (2003) Diacylglycerol Kinase-ζ Localization in Skeletal Muscle Is Regulated by Phosphorylation and Interaction with Syntrophins. Mol. Biol. Cell. 14, 4499–4511

72. Hu, T., and Exton, J. H. (2003) Mechanisms of Regulation of Phospholipase D1 by Protein Kinase Cα*. J Biol Chem. 278, 2348–2355

73. Chen, J.-S., and Exton, J. H. (2004) Regulation of Phospholipase D2 Activity by Protein Kinase Cα*. J. Biol. Chem. 279, 22076–22083

74. Sakane, F., Yamada, K., Imai, S., and Kanoh, H. (1991) Porcine 80-kDa diacylglycerol kinase is a calcium-binding and calcium/phospholipid-dependent enzyme and undergoes calcium-dependent translocation. J Biol Chem. 266, 7096–7100

75. Sakane, F., Imai, S., Yamada, K., and Kanoh, H. (1991) The regulatory role of EF-hand motifs of pig 80K diacylglycerol kinase as assessed using truncation and deletion mutants. Biochem. Biophys. Res. Commun. 181, 1015–1021

76. Campbell, S. T., Franks, C. E., Borne, A. L., Shin, M., Zhang, L., and Hsu, K.-L. (2018) Chemoproteomic discovery of a ritanserin-targeted kinase network mediating apoptotic cell death of lung tumor cells. Mol. Pharmacol. 94, mol.118.113001

77. Takita, T., Konuma, T., Hanazato, M., and Inoue, H. (2011) Diacylglycerol kinase inhibitor R59022-induced autophagy and apoptosis in the neuronal cell line NG108-15. Arch. Biochem. Biophys. 509, 197–201

78. Lavieri, R. R., Scott, S. A., Selvy, P. E., Kim, K., Jadhav, S., Morrison, R. D., Daniels, J. S., Brown, H. A., and Lindsley, C. W. (2010) Design, Synthesis, and Biological Evaluation of Halogenated N-(2-(4-Oxo-1-phenyl-1,3,8-triazaspiro[4.5]decan-8-yl)ethyl)benzamides: Discovery of an Isoform-Selective Small Molecule Phospholipase D2 Inhibitor. J. Med. Chem. 53, 6706–6719

79. Lewis, J. A., Scott, S. A., Lavieri, R., Buck, J. R., Selvy, P. E., Stoops, S. L., Armstrong, M. D., Brown, H. A., and Lindsley, C. W. (2009) Design and synthesis of isoform-selective phospholipase D (PLD) inhibitors. Part I: Impact of alternative halogenated privileged structures for PLD1 specificity. Bioorg. Med. Chem. Lett. 19, 1916–1920

80. Schindelin, J., Arganda-Carreras, I., Frise, E., Kaynig, V., Longair, M., Pietzsch, T., Preibisch, S., Rueden, C., Saalfeld, S., Schmid, B., Tinevez, J.-Y., White, D. J., Hartenstein, V., Eliceiri, K., Tomancak, P., and Cardona, A. (2012) Fiji: an open-source platform for biological-image analysis. Nat. Methods. 9, 676–682

81. Wills, R. C., Pacheco, J., and Hammond, G. R. V. (2021) Quantification of Genetically Encoded Lipid Biosensors, pp. 55–72, Methods in Molecular Biology, 2251, 55–72

82. Peng, D., Vangipuram, M., Wong, J., and Leonetti, M. D. (2024) protoSpaceJAM: an open-source, customizable and web-accessible design platform for CRISPR/Cas insertional knock-in. Nucleic Acids Res. 52, e68–e68

83. Ovesný, M., Křížek, P., Borkovec, J., Švindrych, Z., and Hagen, G. M. (2014) ThunderSTORM: a comprehensive ImageJ plug-in for PALM and STORM data analysis and super-resolution imaging. Bioinformatics. 30, 2389–2390

84. Kim, Y. J., Guzman-Hernandez, M. L., and Balla, T. (2011) A Highly Dynamic ER-Derived Phosphatidylinositol-Synthesizing Organelle Supplies Phosphoinositides to Cellular Membranes. Dev Cell. 21, 813–824

85. Doyle, C. P., Rectenwald, A., Timple, L., and Hammond, G. R. V. (2023) Orthogonal targeting of SAC1 to mitochondria implicates ORP2 as a major player in PM PI4P turnover. bioRxiv. 10.1101/2023.08.28.555163

86. Pemberton, J. G., Kim, Y. J., Humpolickova, J., Eisenreichova, A., Sengupta, N., Toth, D. J., Boura, E., and Balla, T. (2020) Defining the subcellular distribution and metabolic channeling of phosphatidylinositol. J Cell Biology. 219, e201906130

87. Zewe, J. P., Wills, R. C., Sangappa, S., Goulden, B. D., and Hammond, G. R. (2018) SAC1 degrades its lipid substrate PtdIns4P in the endoplasmic reticulum to maintain a steep chemical gradient with donor membranes. Elife. 7, e35588

88. Yanisch-Perron, C., Vieira, J., and Messing, J. (1985) Improved M13 phage cloning vectors and host strains: nucleotide sequences of the M13mpl8 and pUC19 vectors. Gene. 33, 103–119

